# Learning shapes neural geometry in the prefrontal cortex

**DOI:** 10.1101/2023.04.24.538054

**Authors:** Michał J. Wójcik, Jake P. Stroud, Dante Wasmuht, Makoto Kusunoki, Mikiko Kadohisa, Mark J. Buckley, Rui Ponte Costa, Nicholas E. Myers, Laurence T. Hunt, John Duncan, Mark G. Stokes

## Abstract

The relationship between the geometry of neural representations and the task being performed is a central question in neuroscience^1–6^. The primate prefrontal cortex (PFC) is a primary focus of inquiry, as it can encode information with geometries that either rely on past experience^7–13^ or are experience agnostic^3,14–16^. One hypothesis is that PFC representations should evolve with learning^4,17,18^, from a format that supports exploration of all possible task rules to a format that minimises the encoding of task-irrelevant features^4,17,18^ and supports generalisation^7,8^. Here we test this idea by recording neural activity from PFC when learning a new rule (‘XOR rule’) from scratch. We show that PFC representations progress from being high dimensional, nonlinear and randomly mixed to low dimensional and rule selective. Upon generalising the rule to novel stimuli, these representations further evolve into an abstract, stimulus-invariant geometry. These findings reconcile previously conflicting accounts of PFC function by demonstrating how neural representations adapt across distinct stages of learning.

---

Two seemingly discrepant accounts propose that PFC neural activity should track either low-^8–13,19^ or high-dimensional^3,14–16^ representations of the environment. Traditionally, it has been proposed that PFC cells are tuned adaptively to task-relevant information, leading to low-dimensional neural activity^13^. This results in the population displaying structured selectivity patterns, as commonly observed after training on a cognitive task (**Fig. 1a**, *low-dimensional*) ^13^. A contrasting hypothesis suggests that the PFC may rely on high-dimensional, nonlinearly mixed representations of task features to support complex cognition (**Fig. 1a**, *high-dimensional*)^3,14^. According to this notion, the PFC serves as a nonlinear kernel such that when a low-dimensional input is projected onto it, dimensionality expands, and a wide repertoire of responses can be generated^15,16^.

**Figure 1.**
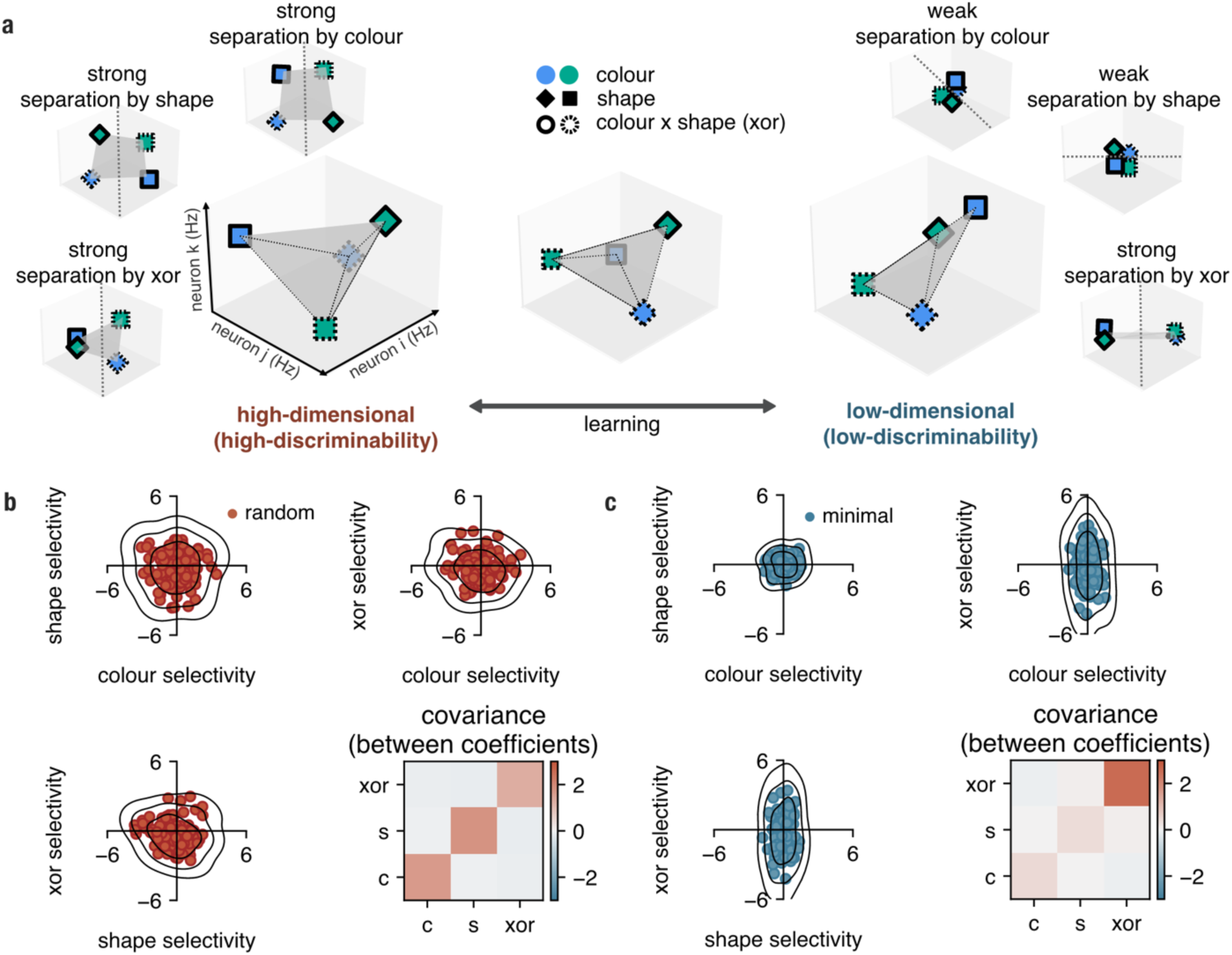
Potential effects of learning on neural geometry in the prefrontal cortex. Learning can reduce or expand neural dimensionality, changing how many linear decoding axes can be implemented on neural firing rates (discriminability). **a**, High-dimensional representations enable high discriminability. A high-dimensional regime allows the strong separation of all task features using three possible readout axes (left), whereas a low-dimensional representation only allows task-relevant features to be strongly separated (right). **b**, Each neuron can be represented as a point in the 3-dimensional selectivity space spanned by colour, shape, and XOR (their interaction). In the random model, selectivity is distributed according to a spherical Gaussian distribution in this space (*Methods, generative models*); the covariance matrix is computed between the selectivity coefficients; zero-mean Gaussian noise (σ = 0.7) was added to each selectivity coefficient to illustrate measurement bias under finite sampling. **c**, Analogous to b but for the minimal model; neurons are strongly selective only for the XOR (interaction between colour and shape), as this is the only feature that is necessary to solve the task.

Recently, it has been proposed that the PFC is capable of transitioning between high- and low-dimensional representations across learning, to accommodate the changing demands of the environment ^4,17,18,20^. For example, early in learning, high-dimensional representations may allow flexible exploration of all possible input–output mappings (“contingencies”) in order to discriminate which task rules are currently relevant ^3,14,16^. This is because a high-dimensional representation allows for a high number of linearly separable task features (**Fig. 1a**). Conversely, once an animal has learnt that only one set of contingencies is relevant, a low-dimensional representation may be used to encode task-relevant features more robustly^4,17–19^. Moreover, these low-dimensional representations may enable generalisation to novel contexts, since aligning new with old representations is likely easier when fewer dimensions must be considered. (**Supp. Fig. 1**)^7,8^. In other words, different stages of learning impose different demands on the neural population. Learning could thus shape neural dimensionality and progressively push neural activity towards different solutions along the trade-off between discriminability and generalisability, i.e., from a high-dimensional regime towards a low-dimensional regime^18,20^.

Here, we tested this idea in two macaque monkeys which learnt an exclusive-or (XOR) rule – a problem that can be solved by a range of representations, from low- to high-dimensional (**Fig. 1a**)^19^. Importantly, we tracked how the dimensionality and geometry of PFC representations changed across multiple training sessions of an XOR rule that was entirely new to the animals at the start of recording (experiment 1) and during subsequent generalisation of this rule to a new stimulus set (experiment 2). We used a classical conditioning paradigm in which the nonlinear combination of the features of two objects presented in succession (XOR) predicted the outcome of the trial. Importantly, the animals were only required to fixate through both experiments^21–24^. Later, we also show that our results hold in a previously collected delayed match-to-sample task^25–27^.

Across two experiments, we found that during early stages of learning, PFC activity was high-dimensional, with individual neurons exhibiting nonlinear and randomly mixed selectivity. As learning progressed, population activity became increasingly low-dimensional, with structured selectivity emerging predominantly for task-relevant variables. When novel combinations of stimuli were introduced, PFC representations reorganised such that new and familiar conditions aligned along a shared axis, enabling the reuse of a common neural code. These findings demonstrate that learning reshapes both the dimensionality and geometry of neural representations in the PFC, promoting low-dimensional and abstract encoding when animals continuously engage in a single task structure.

## Generative models of nonlinear random and minimal selectivity

We first wanted to understand how the geometry of the neural representations could change over the course of learning. We thus explored the geometries produced by two generative models of neural selectivity with different discriminability-generalisability trade-offs^4,17^: (i) a high-dimensional geometry produced by non-linear random mixed selectivity^3,14–16,28^ (high discriminability, low generalisability); and (ii) a low-dimensional geometry produced by structured, minimal selectivity (low discriminability, high generalisability)^10,12,29–33^. The former is a well-established geometry (e.g., inherent in reservoir computing models^15,16^) whereas the latter was inspired by work suggesting that PFC neurons flexibly adapt their selectivity to current task demands^13^. These models make distinct predictions about the distribution of selectivity to the task-relevant variables in a 3-dimensional selectivity space (colour, shape, and their nonlinear interaction, i.e., XOR). We refer to minimal or random selectivity when describing the distribution of selectivity and to low- or high-dimensional geometry when referring to the respective representations generated by these distributions (see *Discussion* for a detailed distinction between dimensionality and selectivity).

In selectivity space, each axis represents a units’ response to one stimulus variable (e.g., high shape selectivity = higher firing rate for square than diamond, **Fig. 1b**). Hypothetically, one could imagine different distributions of variable encoding within this selectivity space. The properties of these distributions are determined by a covariance matrix: the diagonal elements describe the strength of coding of each variable (variance) whereas the off-diagonal entries determine the strength of the relationship between variables (covariance). In our generative models neural firing rates were simply constructed as a linear combination of task variables (colour, shape and XOR inputs represented using one-hot encodings; see *Methods, generative models* for details) with a specified covariance matrix of selectivities to the task variables. In line with previous studies, the high-dimensional model was constructed by allowing selectivity to linear (colour and shape) and nonlinear (XOR) features to be distributed randomly according to a spherical Gaussian distribution (**Fig. 1b**). In contrast, in the minimally structured XOR selectivity model, neurons were only strongly selective for the nonlinear interaction (i.e., the XOR; **Fig. 1c**). One possible way of constructing such a model is considering biologically plausible limits on neural firing rates (minimising net firing rate). In line with this, we derived this model mathematically and demonstrated that it minimises total firing rate activity while maximising task performance (see **Supp. Materials, section 1**). Consistent with this, we found that feedforward networks trained using backpropagation to perform the task while also minimising a metabolic cost term converge to the minimal XOR selectivity model (see **Supp. Fig. 2a-h**; *Methods, optimised feedforward networks*). Please note that alternative mechanisms, such as initialisation or presence of noise, could be also applied to learn a similar minimal selectivity model^10,34^. In line with prior accounts^17,35^, XOR decoding in the low-dimensional model was more robust to noise (**Supp Fig. 2i**) and required fewer units to implement a stable readout (see **Supp. Fig. 2j**).

We established a metric to measure whether neural activity is better described by the random or minimal model. We first fitted a linear model regression to surrogate data generated by both our generative models, in which task variables (colour, shape, and colour x shape (XOR)) were used as predictors of each unit’s firing rate (see *Methods, model section;* for similar analysis see^36,37^; **Supp. Fig. 3a,b)**. Then we measured the average within-model distance between two covariance matrices drawn from the random model (**Supp. Fig. 3c**, red line; *Methods,* *eq. 4*) and the average between-model distance between the covariances drawn from the random and the minimal model (**Supp. Fig. 3d**, blue line, *Methods,* *eq. 5*). To test whether our measure captures learning dynamics, we constructed four artificial populations with varying proportions of random and minimal selectivity. As the proportion of the minimal model in this mixed population increased, it became more dissimilar to the average random model and more similar to the average minimal model (**Supp. Fig. 3c**, black line). A reflected version of these results held true when the minimal model was used as reference (**Supp. Fig. 3d**).

Subsequently, to gain insight into the task geometries that these models generated, we employed an established technique^3,14^ and trained linear decoders to decode all three task variables in both models. As previously suggested^3,14^, a randomly mixed selectivity model yielded a high-dimensional task representation, allowing for all variables, including the nonlinear XOR, to be decoded (**Supp. Fig. 3e**, red; cf. **Fig. 1a**, far left). In contrast, for the minimal model, only the XOR combination of shape and colour, and not shape or colour independently, could be decoded (**Supp. Fig. 3e**, blue; cf. **Fig. 1a**, far right). While both models can perform the task, we expected their representation of the XOR variable to differ fundamentally. On the one hand, the minimal model by design should represent the XOR in a format that generalises over all other task variables. On the other hand, this is not guaranteed in the random model. We verified this intuition using cross-generalised decoding, a method in which a linear decoder is trained to decode a given task variable (e.g., XOR = True vs. XOR = False) on a given set of task conditions (e.g., blue colour) and tested on a different set of task conditions^7^ (e.g., green colour; see *Methods section, cross-generalised decoding*). We found that for the random model, cross-generalised decoding was at chance-level for all task variables (**Supp. Fig. 3f**, red). This is because the random model, by design, exhibits no reliable structure in its representation of variables and therefore these dimensions are represented randomly in relation to each other. In contrast, the minimal model displayed maximal cross-generalised decoding for the XOR variable (**Supp. Fig. 3f,** blue), indicating that it can be decoded regardless of which set of task conditions the decoder is trained and tested on. This suggests that the minimal model is able to represent the XOR in a highly cross-generalisable format (**Fig. 1a**, far right). Consequently, the minimal model also exhibits below-chance cross-generalised decoding for colour and shape (see **Supp. Fig. 3g, h**). Next, we directly compared neural data at each stage of learning to the selectivity (random vs minimal) and neural geometry (low-vs high-dimensional) generated by these models.

## Learning a single task reduces neural dimensionality in the prefrontal cortex

In experiment 1, the animals were trained to combine a colour stimulus (either blue or green) with a subsequently presented shape (either square or diamond) in a nonlinear fashion to predict the reward (the XOR between colour and shape) outcome of the trial (**Fig. 2a,b**). Using a semi-chronic multielectrode system we sequentially recorded 376 neurons from the lateral PFC across both macaques (**Fig. 2c**; see *Methods, data acquisition and pre-processing*). Moving electrodes between sessions ensured that a new sample of neurons was obtained in each session. Importantly, to capture learning dynamics, we started recording from the first session in which the animals were exposed to the task. Experimental sessions were split into four learning stages for each animal separately. Data for each stage was combined across animals. A sliding-window approach was used to utilise all available trials while ensuring an equal number of trials per learning stage; each stage comprised approximately equal numbers of training sessions (see for details see *Methods*, *data acquisition and pre-processing*). Selectivity analyses were run in the time window before the animals received feedback about the outcome of the trial (i.e., reward; for details see *Methods*, *data acquisition and pre-processing*).

**Figure 2.**
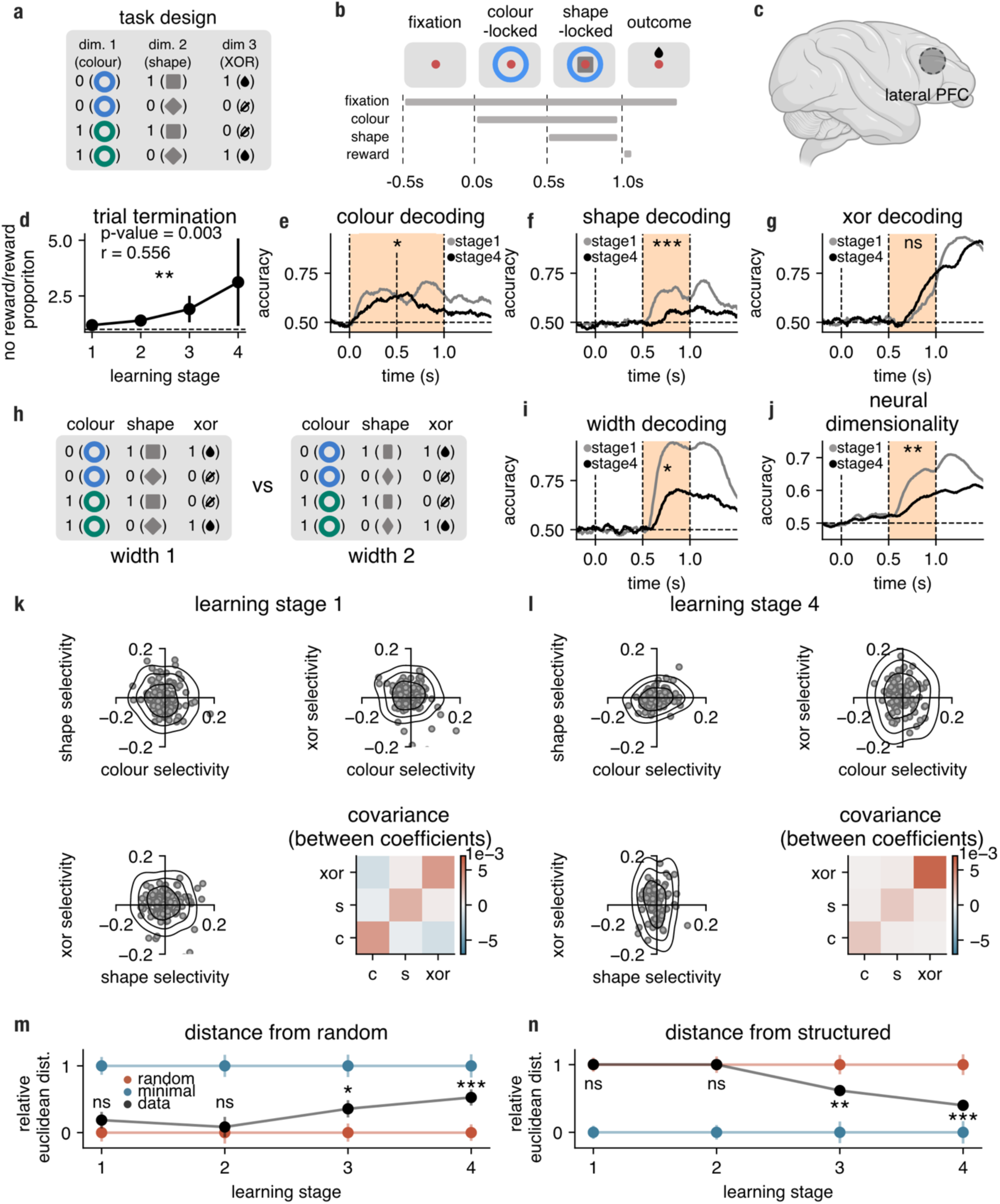
Neural representations in macaque PFC during learning of a single task. **a**, In Experiment 1, animals were incentivised to combine two passively viewed task features (colour and shape) in a non-linear fashion (XOR). For example, blue+square and green+diamond combinations were rewarded, whereas blue+diamond and green+square were not. **b**, Timeline of task events in a single trial. **c**, Neural data was collected from the lateral surface of the prefrontal cortex in two macaque monkeys (see *Methods, data acquisition and pre-processing* for further details). **d**, The tendency of animals to terminate trials in the shape-locked trial period plotted as a ratio of termination numbers in not rewarded and rewarded trials (illustrated as a function of learning). **e-g**, Time resolved linear decoding in stage 1(grey) and stage 4(black) of colour, shape, and XOR, respectively; the pale orange shaded areas denote the time windows in which permutation tests were conducted to assess differences between stage 1 and stage 4 decoding scores; vertical three dashed lines show the onset of the colour, shape and the outcome, respectively. **h,** Schematic of narrow and broad shape trials (width feature). This feature was not predictive of reward. **i**, Temporally resolved linear SVM decoding of width. **j**, Time resolved neural dimensionality as measured by shattering dimensionality (all possible task dichotomies excluding colour, shape, width and XOR). **k**,**l**, Neural selectivities in learning stages 1(k) and 4 (l) computed in the late shape-locked period ([t_400*ms*_, t_500*ms*_], shape-locked). Each point represents the selectivity of one neuron for each of the task variables (colour, shape and XOR). We show all 3 possible pairs of the 3 axes. The contour plots represent a kernel density estimate of the data, with each ring corresponding approximately to 1, 2, and 3 standard deviations from the mean. The right panels show the covariance matrix of the selectivities computed from the data for stage 1 of learning (**k**) and stage 4 of learning (**l**). **m**, Relative Euclidean distance between the covariance matrix of selectivity coefficients from the data ([t_400*ms*_, t_500*ms*_], shape-locked) and the covariance matrix expected from random selectivity (with matched total variance) as a function of learning (*Methods*, *measuring similarity between selectivity distributions*). Red and blue, respectively, error bars show mean (±1 *s. d*. over 1000 randomly drawn models) of the relative Euclidean distance between the covariance matrix of the random, respectively minimal, model (with matched total variance to the data) and the covariance matrix expected from random selectivity; black error bars show standard deviation of relative distance between the observed covariance and random covariance (±1 *s. d*. over 1000 random models). **n**, Same as panel d but we show the relative distance of the observed covariance (([t_400*ms*_, t_500*ms*_], shape-locked) from the covariance matrix expected from minimal selectivity (with matched total variance). All p-values were calculated from permutation tests (***, *p* < 0.01; **, *p* < 0.01; *, *p* < 0.05; †, *p* < 0.1; *n. s*., not significant).

We first assessed learning behaviourally by examining the animals’ tendency to terminate non-rewarded trials before the potential reward onset (shape-locked period, **Fig. 2b**) by breaking fixation (**Fig. 2d**). This was quantified by calculating a trial termination index, i.e. the ratio of terminated non-rewarded trials to terminated rewarded trials, which we tracked across different learning stages (see *Methods, trial termination* for details). Over the course of learning, the animals increasingly differentiated between rewarded and non-rewarded trials (*r* = .56, *p* < 0.01; **Fig. 2d**). We next investigated which learning strategy the animals adopted. One possibility is that they memorised each stimulus combination and its associated outcome (a *flat strategy*; **Supp. Fig. 4a**). Alternatively, they may have used a hierarchical strategy, in which colour served as a first-order policy cue guiding subsequent context-dependent processing of shape (**Supp. Fig. 4b**). To distinguish between these strategies, we analysed switch costs (see *Methods, switch costs* for details), comparing the animals’ ability to terminate non-rewarded trials following a change in colour versus shape from the previous trial (**Supp. Fig. 4c,d**). We found no evidence of shape switch costs (*r* = .01, *p* = 0.41; **Supp. Fig. 4e**), whereas colour switch costs increased over learning (*r* = .37, *p* < 0.05; **Supp. Fig. 4f**) and were significantly greater than shape switch costs (*r* = .36, *p* < 0.05; **Supp. Fig. 4g**). These results suggest that animals increasingly relied on colour as a higher-order cue, consistent with the adoption of a hierarchical learning strategy.

We next applied our linear decoding analyses to the neural recordings to establish whether the emergence of this behavioural strategy was accompanied by changes in the dimensionality of neural representations. At the beginning of training (learning stage 1, grey lines), the animals exhibited a high-dimensional geometry that allowed for colour, shape and XOR to be decoded, reminiscent of the high-dimensional model (**Fig. 2e-g)**. This was especially prominent in the period prior to the reward delivery.

Over the course of learning (stage 1 vs stage 4) we observed a reduction in colour decoding (*p* < 0.05, one-sided) and shape decoding (*p* < 0.001, one-sided) but not XOR decoding (**Fig. 2e-g**, grey vs black lines). The shape stimuli also had a feature that was irrelevant for the prediction of the outcome: width (**Fig. 2h**). Similarly to colour and shape, the decoding of this feature also decreased over learning (*p* < 0.01, one-sided). The reduction of colour and shape coding as well as stable output-relevant feature (XOR) decoding was predicted by a transition from a high-dimensional to a low-dimensional model **(Supp. Fig. 3e**, red vs blue).

We next explicitly tested whether the dimensionality of neural representations changed over learning (as measured with shattering dimensionality; see ref. ^14^ and *Methods, decoding* for details). A neural representation described by three binary input dimensions (colour, shape and width) results in 35 dichotomies (division into two sets of four stimuli) that can be theoretically decoded. We found that the mean decoding accuracy of all dimensions (excluding colour, shape, width and XOR) decreased significantly over learning (*p* < 0.01, one-sided; **Fig. 2j**). Additionally, a principal component analysis computed on condition averages revealed that the proportion of variance explained by the first principal component increased as a function of learning (*M*_*stage*1_ = 0.466 *vs M*_*stage*4_ = 0.581, *p* < 0.05, one-sided; **Supp. Fig. 4h**; for details see *Methods, Principal component analysis*).

We next tested whether changes to neural dimensionality were reflected in changes to selectivity. We fitted a linear regression to our data, just as we did for our generative models (**Supp. Fig. 3a, b**), in which task variables (colour, shape, and colour x shape (XOR)) were used as predictors of each neuron’s firing rate. We then examined how selectivity coefficients changed over learning (**Fig. 2k**, learning stage 1 and **Fig. 2l**, learning stage 4). We compared the covariance structure of these selectivity coefficients (**Fig. 2k, l**, bottom right) to the covariances obtained from the random selectivity model and minimal selectivity model (**Supp. Fig. 3a, b;** covariance). At the beginning of learning (stage 1), PFC cells were randomly distributed in selectivity space resembling the high-dimensional model (*p* = 0.332, one-sided, **Fig. 2m** and *p* = .99, one-sided, **Fig. 2n**). However, in late learning (stage 4), selectivity diverged away from randomly mixed selectivity (*p* < 0.001, one-sided, **Fig. 2m**, compare black and red lines) and converged towards the minimal model (*p* < 0.001, one-sided, **Fig. 2n**, compare black and blue lines).

We replicated a similar pattern of decoding and selectivity results on data sorted by the trial termination index. Here, the behavioural measure served as a performance indicator, and sessions were sorted separately for each animal before being pooled into four pseudopopulations across animals (see *Methods, trial termination measure*; **Supp. Fig. 4i-o**). Furthermore, we also performed a re-analysis of an existing dataset^25–27^ in which recordings were taken from primate ventral and dorsolateral PFC before and after learning a delayed match-to-sample task that was similar in structure to ours (for details see *Methods, existing lPFC dataset*; **Supp. Fig. 5**). We found that learning again pushed neural activity in the PFC towards a minimal regime.

Our findings indicate that neural activity in the PFC shifts between two distinct selectivity regimes as learning progresses. Initially, the PFC maximally expanded the representational space by encoding all available variables. Subsequently, after a combination of task variables that predicted the trial’s outcome were identified, neural activity became increasingly low-dimensional. Such changes to neural dimensionality can be associated with simultaneous changes in neural geometry, potentially supporting more abstract representations (**Supp. Fig. 3f**). To test this, we employed cross-generalised decoding and, consistent with our generative models, observed a strong increase in cross-generalised XOR decoding across learning (*p* < 0.001, one-sided; **Supp. Fig. 4p**). Although this effect is consistent with abstraction, it could also reflect motor-preparation or reward-prediction signals injected into the PFC in the rewarded XOR condition (XOR==True). To directly assess whether an abstract, task-relevant dimension emerges independently of such signals, we conducted experiment 2, introducing a new task instance that preserved the same underlying structure.

## Learning a single task structure promotes abstract neural geometry in the PFC

In experiment 2, we introduced a new colour pair (*stimulus set*; **Fig. 3a**) that followed the same shape–outcome associations as the previous colour pair (*context*, **Fig. 3b**). Similar to experiment 1, we recorded neural activity from the very first session in which the animals were exposed to the new stimulus set and divided the experimental sessions into four distinct learning stages. To explore learning-induced changes to neural dimensionality, we again employed linear decoding. We next compared these metrics to changes in the structure of neural selectivities. During the colour-locked period, PFC activity could now represent three distinct variables (**Supp. Fig. 6a**): context (i.e., the colour indicating the relevant shape–outcome rule), stimulus set (set 1 vs. set 2), and the nonlinear interaction between context and stimulus set (XOR 2). Using these three variables we constructed selectivity models analogous to those in **Fig. 2k,l** (**Supp. Fig. 6b-c**). Critically, in this design, context was the only task-relevant variable, whereas both the stimulus set and its nonlinear interaction with context (XOR 2) were irrelevant. We predicted that, as in experiment 1, PFC activity would transition from high-dimensional and nonlinearly mixed to low-dimensional and structured, with cells becoming increasingly selective for the task-relevant variable **(Supp. Fig. 6a)**. Additionally, randomly interleaving stimulus set 1 trials and stimulus set 2 trials allowed us to test whether a shared neural representation would be used for both stimulus sets. Importantly, none of the analyses performed on activity during the colour-locked period were confounded by motor preparation or reward prediction, as the shape information required to predict reward had not yet been presented.

**Figure 3.**
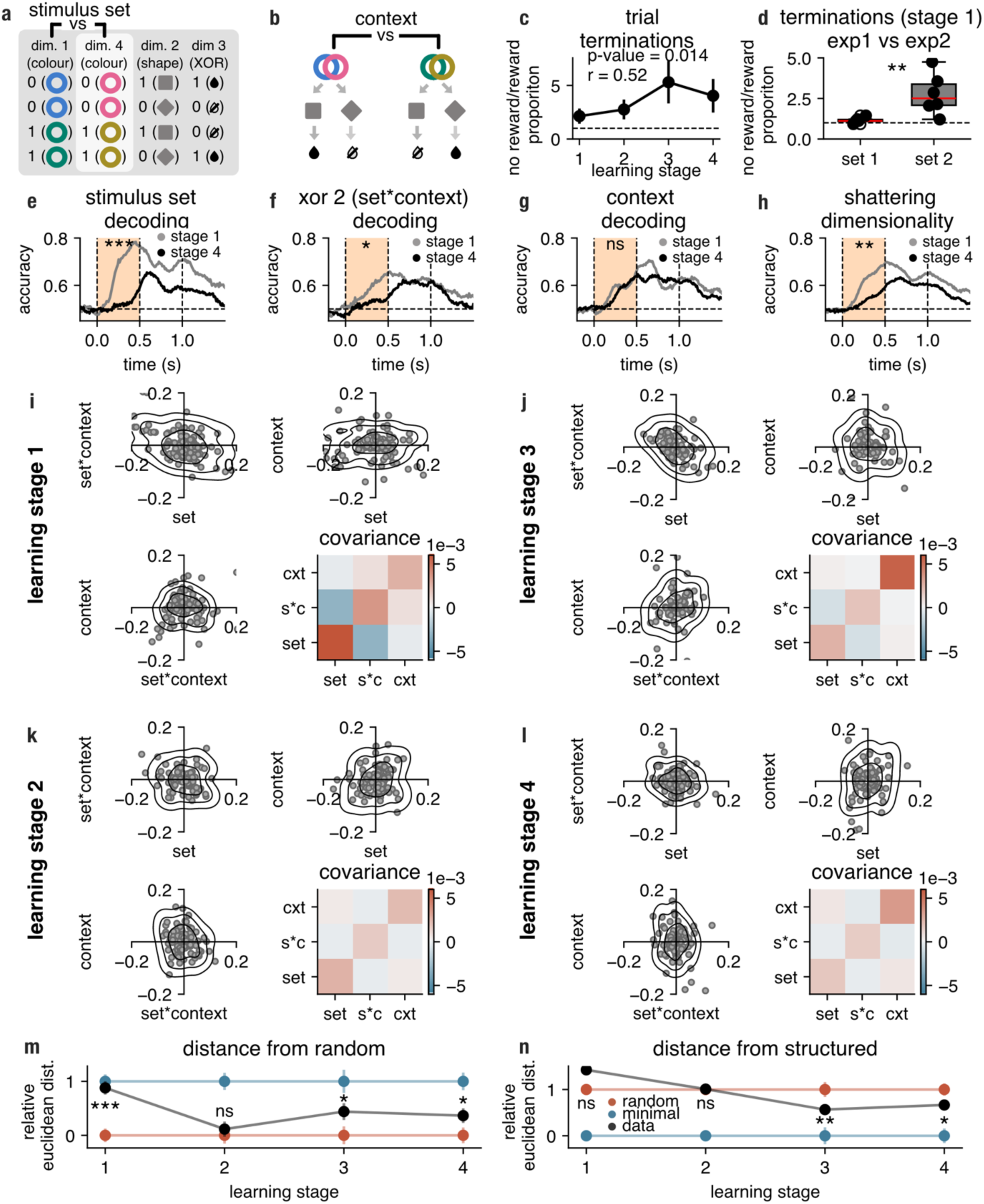
Neural representations in macaque PFC during learning of two instances of the same task structure. **a, b**, New colours (pink and khaki, stimulus set 2) were introduced in experiment 2, sharing the same shape–reward mapping as the learned colours (blue and green, stimulus set 1). **c**, Analogous to Fig. 2a but computed for stimulus set 2 trials. **d**, Comparison of the trial terminations observed in the first 3 sessions (per animal) in experiment 1 (when only stimulus set 1 was presented) and 3 first sessions (per animal) in experiment 2 for stimuli set 2 trials only. **e-g**, Temporally resolved linear SVM decoding stimulus set, XOR 2 (set*context) and context; the pale orange shaded areas denote the time windows in which permutation tests were conducted to assess differences between stage 1 and stage 4 decoding scores. Horizontal dotted lines represent chance-level decoding whereas vertical dotted lines indicate the onset of the colour, shape and the trial outcome. **h**, Time resolved neural dimensionality as measured by shattering dimensionality (mean over all 3 possible task dichotomies in the colour-locked period, i.e., mean over panels e-g) in stage 1 and stage 4. **i**-**l**, Neural selectivities in learning stages 1-4 computed in the colour-locked period ([t_200*ms*_, t_500*ms*_]). Each point represents the selectivity of one neuron for each of the task variables (set, XOR 2 and context). We show all 3 possible pairs of the 3 axes. The contour plots represent a kernel density estimate of the data, with each ring corresponding approximately to 1, 2, and 3 standard deviations from the mean. The right panels show the covariance matrix of the selectivities. **m**, **n**, Analogous to Fig. 2m,n; compares set, XOR 2 and context selectivities ([t_200*ms*_, t_500*ms*_], colour-locked) to idealised random and minimal models as a function of learning. All p-values were calculated from permutation tests (***, *p* < 0.01; **, *p* < 0.01; *, *p* < 0.05; †, *p* < 0.1; *n. s*., not significant).

We first examined the propensity of animals to terminate trials when the new stimulus set indicated a lack of reward, similar to the patterns observed in experiment 1. Over the course of learning with stimulus set 2, animals increasingly terminated non-rewarded trials more frequently than rewarded ones (*r* = .52, *p* < 0.05, **Fig. 3c**). Additionally, a facilitation effect was observed when comparing the first three sessions of experiment 2 (stimulus set 2 only) to the first three sessions of experiment 1 (stimulus set 1). Specifically, animals demonstrated significantly more adaptive trial termination early in the learning process with stimulus set 2 compared to stimulus set 1 (*p* < 0.01, **Fig. 3d**). This early behavioural benefit may be attributed to the utilisation of the previously acquired task representation as a scaffold. Subsequently, we thus investigated how the neural representations of both stimulus sets interacted throughout the learning process.

Similarly to experiment 1, we hypothesised that the PFC would strongly represent all three variables—stimulus set, context, and XOR 2 (i.e., context × stimulus set)—at the beginning of learning, consistent with a high-dimensional coding regime (left, **Supp. Fig. 6a**), and would then progressively transition to a low-dimensional representation in which most neurons are selective for context, the only task-relevant variable (right, **Supp. Fig. 6b**). To test this hypothesis, we first employed linear decoding (**Fig. 3e-g**).

We found that both stimulus set decoding (*p* < 0.001, one-sided) and XOR 2 decoding (*p* < 0.05, one-sided) significantly decreased over learning, whereas context decoding remained stable across learning stages, mirroring the decoding results from experiment 1. Furthermore, we observed a significant reduction in shattering dimensionality during the colour-locked period as learning progressed (*p* < 0.01, one-sided). No learning-related changes were detected for shape or XOR representations (**Supp. Fig. 7a, b**). Notably, width decoding further declined with learning in experiment 2 (*p* < 0.05, one-sided; **Supp. Fig. 7c**).

To determine whether these dimensionality changes were reflected in the structure of neural selectivity, we analysed selectivity for the three task variables (context, set, and XOR 2) at each learning stage and compared the population profiles to idealised random and minimal models (**Supp. Fig. 6b** and **c**, respectively). In stage 1, neural selectivity significantly diverged from the random model and was inconsistent with the minimal model (*p* < 0.001, one-sided; **Fig. 3m**), driven by a strong preference for the stimulus set variable (stage 1; **Fig. 3i**). We interpret this as a novelty effect, with different neural responses for novel vs. familiar stimuli. In stage 2 (**Fig. 3k**), PFC selectivity shifted away from this initial set-selective regime and converged toward a profile consistent with randomly mixed selectivity (*p* = 0.825, one-sided; stage 2; **Fig. 3m**). From there, it gradually transitioned toward the minimal regime in stages 3 and 4 (**Fig. 3j,l**), reflecting increasingly structured and task-relevant coding (*stage* 3: *p* < 0.05, one-sided, *stage* 4: *p* < 0.05, one-sided, **Fig. 3m**; *stage* 3: *p* < 0.01, one-sided, *stage* 4: *p* < 0.05, one-sided, **Fig. 3n**). Together with experiment 1, these results identify random mixed selectivity as a critical stage in early learning, even when new information must first be integrated with existing representations. Subsequently, whether learning a completely novel task (experiment 1) or a task related to prior knowledge (experiment 2), the PFC progressively converged towards minimal selectivity.

We next examined whether the observed changes in dimensionality were accompanied by changes in neural geometry. Specifically, we assessed how task variables—context, shape, and XOR—were coded across stimulus sets. For example, we asked whether the decision boundary separating the two contexts in stimulus set 1 (blue vs. green) was the same as that in stimulus set 2 (pink vs. khaki; **Fig. 3b**). We found that at the beginning of learning (stage 1) PFC cells that were selective to context in stimulus set 1 tended to be not selective to context in stimulus 2 (*r* = −.15, *p* > 0.05, one-sided, **Fig. 4a**, left). After learning, in stage 4, the same neurons exhibited context selectivity that generalised across stimulus sets (*r* = .37, *p* < 0.001, one-sided, **Fig. 4a**, right). Consistent with this, correlations between single-neuron context selectivity across colour pairs increased progressively over learning (*p* < 0.01, one-sided, **Fig. 4b**). These changes at the single-neuron level were mirrored by a reorganisation of population geometry. To quantify this, we trained a linear SVM to decode context in stimulus set 1 and evaluated its performance on stimulus set 2, and vice versa (see *Methods, cross-stimulus set generalisation*; **Fig. 4c-d**). At the end of learning (stage 4), context decoding robustly generalised across stimulus sets, indicating that a common decision boundary was used regardless of the stimulus set presented (*p* < 0.001, one-sided; **Fig. 4c**, right). This geometry emerged with learning, reflected in an increase in cross-set generalised decoding from stage 1 to stage 4. (*p* < 0.05, one-sided; **Fig. 4d**, grey, ‘cross-decoding’). Note that context was equally decodable in both early and late learning, indicating that the observed change in geometry was not driven by a change in the presence or absence of context information (*p* > 0.05, two-sided; **Fig. 4d**, black, ‘decoding’). These generalisation effects were robust to how learning was discretised into stages (**Supp. Fig. 7e**). Moreover, context cross-set generalisation reached the ceiling level of simple context decoding after learning (**Supp. Fig. 7f**), indicating that an abstract, stimulus-invariant geometry came to dominate the neural representation of context.

**Figure 4.**
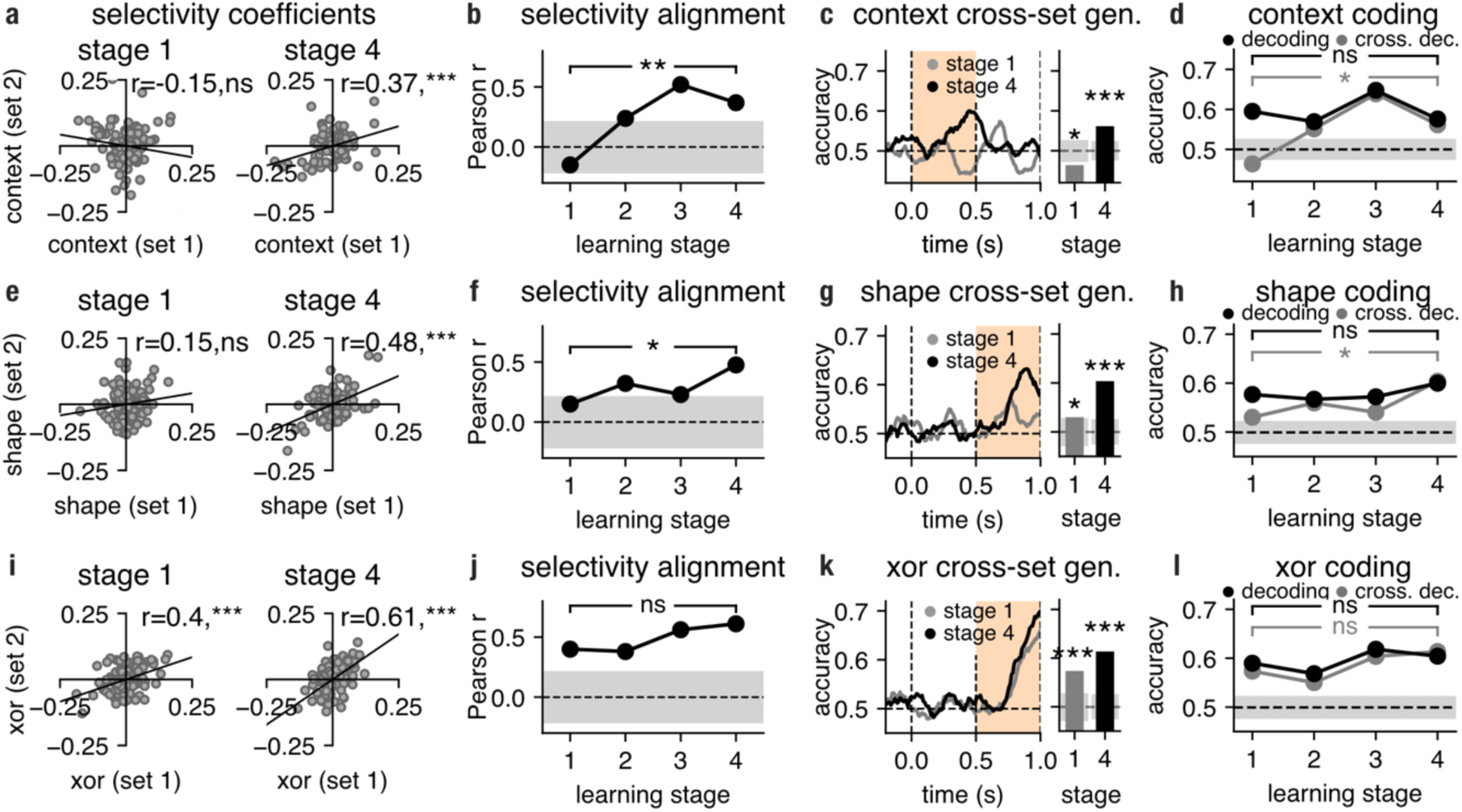
The PFC aligns new and old task representations as a function of learning. **a**, Selectivity of PFC neurons in stage 1 and stage 4 for context in set 1 and set 2 in the colour-locked period. The line of best fit is shown in black. **b**, Correlation between selectivity coefficients (computed in the colour-locked period, [t_0*ms*_, t_500*ms*_]) for context (context 1 vs context 2) in stimulus set 1 and stimulus set 2 in late the colour-locked period plotted as a function of learning (cf. panel a). **c**, Temporally resolved cross-generalised decoding of context (trained on set 1 and tested on set 2, and vice-versa); pale orange shading indicates when the decoding and selectivity analyses in panels a-d were performed. Horizontal dotted lines represent chance-level decoding whereas vertical dotted lines indicate the onset of the colour, shape and the trial outcome. The bar plot represents the average cross-set generalisation score of stage 1 and stage 4 compared against a null distribution obtained after shuffling trial labels (grey-shaded areas). **d**, Decoding (black) and cross-generalised decoding (grey) of context as a function of learning in the colour-locked period (pale orange area in panel c). The grey shaded areas indicate combined chance-level cross-set gen. decoding. **e-l**, Analogous to a-d but for shape and XOR coding and selectivity. All p-values were calculated from permutation tests (***, p < 0.01; **, p <0.01; *, p < 0.05; †, *p* < 0.01; n.s., not significant).

Stimulus set-invariant representations were also observed for shape and XOR. For shape, PFC neurons showed no significant alignment of selectivity across stimulus sets at stage 1 (*r* = 0.15, *p* > 0.05, one-sided) but exhibited significant alignment after learning at stage 4 (*r* = 0.48, *p* < 0.001, one-sided; **Fig. 4e**). This increase in correlation was learning-dependent (*p* < 0.05, one-sided; **Fig. 4f**). Cross-set generalised decoding of shape was significant both early (stage 1; *p* < 0.05, one-sided) and late (stage 4; *p* < 0.001, one-sided; **Fig. 4g**). Although, shape decoding did not change with learning (*p* > 0.05, two-sided; **Fig. 4h**, black), cross-set generalised shape decoding increased over learning (*p* < 0.05, one-sided; **Fig. 4h**, grey). These effects were robust across learning discretisations (**Supp. Fig. 7g**). As for context, cross-stimulus-set generalisation approached the ceiling of simple shape decoding after learning, indicating dominance of a stimulus set-invariant representational geometry (**Supp. Fig. 7h**). For XOR, selectivity alignment and cross -set generalised decoding were already high at stage 1 and remained high at stage 4 (**Fig. 4i–k**), with learning providing no further improvement in these measures (**Fig. 4j, l**). These results were robust across learning discretisations (**Supp. Fig. 7i**), and cross-set generalised XOR decoding remained near ceiling throughout learning (**Supp. Fig. 7j**).

Together these results show that even when new information fits an existing task schema, the PFC traverses a representational trajectory from high-dimensional, mixed coding toward low-dimensional, task-relevant abstraction. Critically, abstraction emerges via progressive alignment of neural geometry across stimulus sets, with some variables (XOR) generalising almost instantly, while others (context, shape) require representational reorganisation.

## Discussion

The prefrontal cortex has the capacity to generate both low-dimensional and high-dimensional representations, each of which presents a unique trade-off between generalisability and discriminability. However, the conditions under which each regime is employed currently remain unclear. Our study investigated how the dimensionality and geometry of neural activity changed over learning. We observed that, as learning progressed, neural activity in the PFC transitioned from being high-dimensional with high-discriminability to being low-dimensional and abstract. This transition in the representational strategy was accompanied by a change patterns of single cell selectivity, from random and nonlinearly mixed towards minimal and structured. The structured representations that emerged during learning then supported the generalisation of the learned rule to a novel stimulus set.^18,20^

We found that the PFC transitioned from a high- to a low-dimensional regime over multiple days of exposure to a complex task. This corroborates the findings of Hirokawa et al.,^12^ who found that neural activity covaried with behaviourally relevant variables, thus occupying a low-dimensional manifold. On the other hand, some studies have suggested that an increase in neural dimensionality is predictive of performance^14^. It is possible that the structure of the task and training provides an explanation for these contrasting findings^18^. In our study, recordings were initiated from the onset of task training, spanning a period of five weeks, with animals experiencing the entire task structure from the first session. In contrast, many other investigations into the primate PFC’s involvement in complex cognitive tasks train animals in a fashion that decomposes the task into multiple subcomponents and either builds up task knowledge across training^14^ or presents them in a serial (block-wise) manner^7,38^. Additionally, whereas in our study one source of information (width) was always irrelevant, in other studies information becomes periodically relevant and irrelevant across multiple blocks, which may promote encoding of currently irrelevant information^37^. These training differences may promote information encoding in a high-dimensional manner that favours discriminability over abstraction.

The acquisition of a low-dimensional representation following the learning of a single XOR rule raises the question of which regime the PFC might adopt when confronted with more complex tasks. Although XOR operations necessitate nonlinear integration and abstraction from sensory input, they can be reduced to simple stimulus-response pairings once the rule has been learnt or when a memory-based strategy is utilised. However, some tasks are more difficult to decompose, as they require switching between multiple orthogonal or conflicting subtasks. It has been suggested^18,20^ that in conditions requiring the performing of multiple tasks in series, a high-dimensional representation could be employed by the PFC in order to maximise flexibility and prevent interference.

Our analytical approach aligns with a growing body of work that emphasises the connection between population structure and neural coding^39^. Specifically, we investigated how diverse forms of single-cell selectivity contribute to the geometry and dimensionality of population-level representations. Our results reveal a negative correlation between neural dimensionality and the emergence of structured selectivity. However, the direction of this relationship may be more nuanced and task-dependent. For instance, a highly structured population with pure selectivity for multiple variables might exhibit higher dimensionality than a randomly mixed population responding to fewer variables, even if the latter is nonlinear.

Our findings suggest that the PFC could employ a multi-phase learning strategy, involving distinct temporal dynamics. Initially, novel tasks could be solved via flexible, reservoir-like dynamics, bypassing the need for immediate synaptic plasticity. As training progresses over longer timescales, the PFC could gradually refine its local connectivity, optimising for performance. This dual-phase approach enables both rapid adaptation and efficient resource allocation, echoing models of the cerebellum-motor cortex interactions, where the cerebellum rapidly drives cortical activity through input control^40^. Similarly, an external region could modulate the PFC’s activity on shorter timescales, enabling flexible high-dimensional representations. Over time, the PFC’s intrinsic circuitry would consolidate these representations and assume direct task control. Future research could explore the geometry of task representations acquired at different learning stages and the critical role of synaptic plasticity in this process.

The implicit assumption in many experimental paradigms is that the animals are presented with tasks as tabula rasa, devoid of prior knowledge or training. However, it is unlikely that the animal’s entire experimental history, including life experience, is irrelevant to a given task. Our second experiment allowed us to address this issue and explicitly explore the interactions between already learnt and new information. In line with previous predictions^7,17^, we found that when a new task instance is added, both the new and old instances were rapidly aligned to common axes and sensory differences between them were collapsed. Notably, different task motifs exhibited distinct generalisation timescales: the XOR representation generalised early in learning, while the context motif required weeks of training to generalise. This disparity likely reflects differences in their structural composition. The XOR rule’s use of identical shapes across tasks likely facilitated rapid alignment, leveraging existing neural encoding schemes. In contrast, the context motif’s novel colours necessitated additional encoding and adaptation in the prefrontal cortex, slowing generalisation. This suggest that PFC’s representational alignment is modulated by the degree of overlap between prior and novel stimuli. Shared features could thus promote efficient transfer of learned representations, while novel features could impose additional encoding demands.

It is perhaps surprising that, given the key role of the PFC in the development and acquisition of structured knowledge, only a few studies have investigated how the structure of PFC representations changes during several training days of an entirely novel task ^41,42^. By tracking changes in neural activity across learning, it is possible to identify the biological principles that are required to produce representations supporting higher cognitive functions^43^. Future experiments should extend this paradigm, to track changes in learning even more complex and naturalistic tasks^44^; those that have a compositional structure^45,46^; the influence of different learning curricula^47^; and how these representations change within the same individual neurons as opposed to pseudo populations^48–50^.

## Author contributions

M.J.W., D.F.W, J.D., and M.G.S. conceived the study. Ma.K and M.J.W. coded the experimental procedure. J.D. led the experimental recordings, Ma.K., and Mi.K. performed the experimental recordings; M.J.W. and Mi.K performed data pre-processing. J.P.S., M.J.W., and M.G.S. developed the theoretical framework; J.P.S. performed analytical derivations and constructed the optimized networks. J.P.S. and M.J.W. constructed the generative models and designed the analysis methods. M.J.W. performed the data analysis and produced the figures. J.P.S., L.T.H, N.E.M and D.F.W. supervised and reviewed the data analysis. M.J.W., J.P.S., L.T.H, N.E.M, D.F.W. and J.D. interpreted the results and wrote the manuscript. All authors revised the final manuscript.

## Acknowledgements

This work was funded by the Wellcome Trust (Sir Henry Wellcome Postdoctoral Fellowship to J.S. (215909/Z/19/Z) and Sir Henry Dale Fellowship L.T.H (208789/Z/17/Z), and award 101092/Z/13/Z to M.Ku., M.B., and J.D.), the Strategic Longer and Larger grant (awarded to L.T.H; BB/W003392/1), the Medical Research Council UK Program (MC_UU_00030/7 to M.Ku., M.B., and J.D.), the Biotechnology and Biological Sciences Research Council (award BB/M010732/1 to M.G.S.), Clarendon Fund and Saven European Scholarship (M.J.W), and the James S. Mc-Donnell Foundation (award 220020405 to M.G.S.). Figure 2c was created with BioRender.com. For the purpose of open access, the authors have applied a CC-BY public copyright licence to any author accepted manuscript version arising from this submission. We thank Emilia Piwek and Christopher Summerfield for useful feedback and detailed comments on the manuscript.

## Competing interests statement

The authors declare no competing interests.

## Data availability, and code availability

All code for this study was custom written in Python using the NumPy, SciPy, Matplotlib, Scikit-learn, and TensorFlow libraries. The code repository is publicly available on [GitHub](https://github.com/m-j-wojcik/pfc_learning). The dataset supporting the findings of this study is accessible through [Dryad](https://datadryad.org/stash/dataset/doi:10.5061/dryad.c2fqz61kb). For any additional details, please contact the corresponding author.

## Methods

### Data and task

#### Animals and task

Two adult male rhesus macaques, monkey 1 and monkey 2, were trained in this study. The experiments were conducted in line with the Animals (Scientific Procedures) Act 1986 of the UK and licensed by a Home Office Project License obtained after review by Oxford University’s Animal Care and Ethical Review committee. The procedures followed the standards set out in the European Community for the care and use of laboratory animals (EUVD, European Union directive 86/609/EEC). The animals were seated in a sound- and lighting-attenuated experimental booth. Their heads were restrained and faced a 19-inch screen. The centre of the screen was aligned with a neutral eye position. The animals performed a passive object-association task (**Fig. 2a-b**). Importantly, the animals were accustomed to an experimental setting but had no previous exposure to the task or stimuli introduced in this protocol. Neural recordings were collected from the first session as one of the main aims of the study was to capture learning dynamics. In the first experiment, the animals were presented with a colour and a shape, a nonlinear combination of which predicted reward (**Fig. 2a-b**). In experiment 2, a second set of stimuli was additionally introduced to test whether the rule learnt in the first experiment cross-generalised to the new sensory domain (**Fig. 3a**). The colours used in the coloured circles were designed in the CIELab colour space^51^. The L parameter (luminance) was kept constant which ensured that the stimuli were approximately isoluminant; parameters a and b varied with regard to valence but not value which resulted in a circular colour representation^51^. As colours were randomly assigned to conditions for each animal, this circular representation ensured that regardless of which colour pair was assigned to which XOR mapping, the initial colour similarity/dissimilarity within colour pair was kept constant. Additionally, in both experiments, the second object had two features: one relevant for reward prediction (shape, **Fig. 2a**) and one irrelevant (width, **Fig. 2h**) (for the duration and sequence in which stimuli were presented see **Fig. 2b**) The trial sequence was randomised. All trials with fixation errors were excluded. The dataset contained on average 237.9 (*SD* = 23.9) and 104.8 (*SD* = 2.3) trials for each of the 8 conditions in experiment 1, and 101.0 (*SD* = 18.6) and 54 (*SD* = 1.1) trials per each of the 16 conditions in experiment 2, for monkey 1 and monkey 2, respectively.

#### Data acquisition and pre-processing

Before the start of the experimental protocol, a titanium head holder with two recording chambers was placed and fixed with stainless steel screws in each animal. The frontal recording chambers were implanted over the lateral prefrontal cortex (lPFC) of the right hemisphere in both animals. Data from a second chamber targeting inferotemporal cortex in the right hemisphere are not considered here. A craniotomy was made beneath each chamber to enable electrophysiological recording. Recording locations for each animal are shown in **Supplementary Figure 8**. Surgical procedures were carried out under general anaesthesia and were aseptic. A semi-chronic micro-drive system (SC-96, Gray Matter Research) with 1.5 mm interelectrode spacing, interfaced to a multichannel data acquisition system (Cerebus System, Blackrock Microsystems) was used for frontal recordings. Data were recorded over a total of 25 daily sessions in each monkey (monkey 1: 17 sessions in experiment 1 and 8 sessions in experiment 2; monkey 2: 10 sessions in experiment 1 and 15 sessions in experiment 2). The switch to experiment 2 was made after the animal showed a robust reward prediction signal. Notably, electrodes were manually advanced by a minimum of 62.5 μ*m* before every session to ensure that activity from new cells was recorded. Neural activity was amplified, filtered (300 *Hz* − 10 *kHz*), and stored for offline pre-processing and analysis. Cluster separation was applied (valley seeking algorithm), and the binary spike train was smoothed using a Gaussian window (σ = 50*ms*). We collected spiking activity from 146 and 230 neurons in experiment 1 and from 205 and 151 neurons in experiment 2, for monkey 1 and monkey 2, respectively. Only cells sampled from the ventral and dorsal lateral frontal cortex were included in the data (**Supp. Fig. 8**). No neurons were excluded based on their selectivity profiles. Importantly, as the focus of this study was to track how learning influenced neural geometry and not the magnitude of firing (e.g., repetition suppression effects), we z-scored firing rates of each neuron across the whole session. The obtained firing rate data were then epoched from 200 *ms* before to 1200 *ms* after the colour onset. Next, the full set of sessions in each animal were divided up into four learning stages and then sessions in each stage were pooled across animals, e.g., the first learning stage was comprised of first 5 sessions from monkey 1 and first 3 sessions from monkey 2. We found that four learning stages were sufficient to capture learning-induced effects. To assess whether the choice of learning-stage discretisation influenced the results in Fig. 4, we repeated the analyses using 3, 4, 5, and 6 learning stages. For all discretisations, each stage was required to include a minimum of six sessions, ensuring adequate statistical power. For the three- and four-stage schemes, sessions were grouped into approximately equal blocks. For the five- and six-stage schemes, a sliding-window approach was used (e.g., stage 1 = sessions 1–6, stage 2 = sessions 5–10) to maintain comparable neuron counts and hence statistical power across discretisations. All analyses were implemented in Python using custom-written code and run on combined data (monkey 1 and 2). Two types of analyses were used in this study: (1) timepoint-resolved, where a specific method was applied to every time point in the epoch to track how representations evolved in trial time, and (2) time-averaged, where a method was run on time-averaged data (e.g., [t_0*ms*_, t_500*ms*_], colour-locked or shape-locked) in the time window preceding the shape display or trial outcome. In the former time window, we examined the neural geometry when only the colour information is known, whereas just before outcome onset, we examined whether neural geometry reflected the colour and the shape and their combination (XOR) before the animals received feedback about the value of the trial.

#### Adaptive trial termination and switch costs

To assess learning, we measured the proportion of trial terminations through fixation breaking in both rewarded and non-rewarded trials. Specifically, for each session, we counted the number of trial terminations in rewarded and non-rewarded trials when fixation breaking occurred during the shape-locked period (**Fig. 2b**), a phase where all necessary information for outcome prediction is available but the reward is not yet delivered. These counts were then normalised by dividing by the total number of fixation errors recorded in the session. The adaptive trial termination measure was computed by dividing the normalised non-reward trial count by the normalised rewarded trial count for each session separately. We next divided sessions into four learning stages and fitted a linear regression model with learning stage as the predictor of the adaptive trial termination. To estimate the p-value, we employed a permutation approach, randomising the session-to-learning stage association (*n* = 10,000 permutations). Switch costs were computed by comparing correct trial termination counts between colour-switch and colour-repeat trials; to isolate colour-specific effects, shape-switch costs were subtracted from colour-switch costs.

#### Existing lPFC dataset

We also used an existing dataset of electrophysiological recordings^25^ which have been described in detail previously^26,27^. In brief, neural activity was recorded from the ventral and dorsal lateral PFC (similar to the areas targeted in this study) in four rhesus monkeys who performed a feature match-to-sample task. More specifically, the animals were required to report after a delay period whether the shape of the first stimulus was the same as the shape of the second stimulus. Note that a match/no-match rule is equivalent to an XOR rule. Importantly, neural activity was recorded before the animals were exposed to the task rule (passive viewing) and after they had learned the rule. As both correct match and correct no-match trials were rewarded, the match/no-match signal was not confounded with a reward prediction signal. To test whether neural activity was pushed towards a minimal regime in such experimental conditions we employed the same decoding and selectivity measures as used in the analysis of our dataset (see **Fig. 2**). We examined neural data averaged across the presentation of the second stimulus and the subsequent delay period ([t_0*ms*_, t_2000*ms*_]; stimulus 2-locked). Furthermore, neural activity was analysed for all stimulus pairs combined. For the 8 stimuli, we paired them into 4 sets of pairs and performed our analyses separately on each pair of stimuli (and averaged results over all 4 pairs) so that chance decoding was the same as in our dataset (i.e., 50%).

## Models

### Multiple linear regression

We can model the firing rate ***r*** of a neuron (either from our generative models or our data) at a given time point as a linear combination of the three main task variables: colour, shape and the interaction between colour and shape (the XOR term):

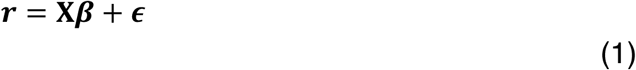

where ***r*** is a vector of 1x K dimensionality containing the time-averaged firing rates for K trials; **X** is the design matrix of dimensionality K x D where rows correspond to the K trials and columns correspond to the value of the D task variables such as colour, shape and XOR (D = 3) in each trial. **β** is a D-by-1 vector populated with the coefficients for each of the task variable estimated for the *n*th neuron. Finally, ε contains K residuals. The **β** vector specifies the coordinates of the *n*th neuron in the selectivity space spanned by D task variables (**Fig. 1b,c** and **Fig. 2k,l**). That is, every neuron can be represented as a point in a space where each axis corresponds to the cell’s selectivity for a task variable. An equivalent linear model was employed to characterise the firing rate for neurons in experiment 2 as a linear combination of the three variables context, stimulus set, and the non-linear mixture of context and stimulus set (XOR2).

### Generative models

Neural selectivity can be defined by the matrix ***S***_*data*_ = (S_*nd*_)_1≤*n*≤*N*,1≤*d*≤*D*_, where each row *n* corresponds to a unit and each column *d* contains the regression coefficient for one task variable. In experiment 1, these variables were colour, shape, and their interaction (colour × shape; XOR); in experiment 2, they were stimulus set, the interaction between stimulus set and context (XOR 2), and context. This cloud of points is then centred by removing the mean (∑_*n*_ *S*_*nd*_ = 0, for each of the *D* task variables). Here, we explored two types of selectivity distributions and their representational properties. Firstly, we examined a random mixed selectivity model in which selectivities are captured by a spherical multivariate Gaussian distribution ***S***_*random*_ ∼ *N*_*d*_(**0**, σ^2^**I**_***d***_). In such a model, all variables can be decoded equally well from the population resulting in a high-dimensional representation and there is poor cross-generalisation between variables. The second selectivity model we examine results from a system performing the task while being constrained to exhibit low overall firing rates (i.e., a form of metabolic cost). We derived analytically that maximising XOR (experiment 1) or the context (experiment 2) decodability while minimising such a metabolic cost results in units being selective only to the task-relevant variable and having no selectivity to the linear terms (colour or shape; **Supp. Materials Section 1**). A matrix describing the selectivity of such a population can be thus formulated as ***S***_*minimail*_ ∼ *N*_*d*_[**2**, σ^2^diag(0,0,1)’, where the covariance matrix is an diagonal matrix with two first diagonal terms equal to zero and the third equal to one. We call this the minimal model. Importantly, to allow comparisons between the observed selectivity and model selectivity (minimal or random), the generative models were constructed using parameters derived from the data. Specifically, we used the mean value of diagonal entries of the covariance matrix 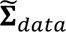 estimated from ***S***_*data*_ to set the value of the variance parameter σ^2^ in both ***S***_*random*_ and ***S***_*minimal*_ (note that this ensures that both models have the same total variance). Furthermore, we showed that multiple linear regression was able to recover the underlying minimal and random models from artificially generated firing rates under various levels of noise (**Supp. Fig. 9**). To mimic measurement variability under finite sampling, we added zero-mean Gaussian noise (σ = 0.7) to the generated selectivity coefficients (**Fig. 1**, **Supp. Fig. 6**). Quantitative comparisons were performed between the observed neural selectivity coefficients and the corresponding idealised noise-free models (**Supp. Fig. 3a, b**).

### Optimised feedforward networks

We used *N* = 400 units in these networks and their firing rates were described by eq. 1 with σ = 2. The output ***z*** of these networks was given by a softmax readout

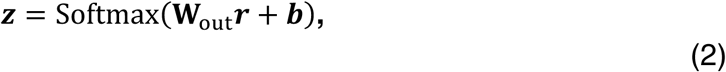

where **W**_out_ are the two sets of readout weights (connecting the hidden layer to the readout unit 1 (XOR == 0) and weights connecting the hidden layer to readout unit 2 (XOR == 1)) and ***b*** is the readout bias. We optimized these networks with back-propagation using a canonical cross-entropy cost function

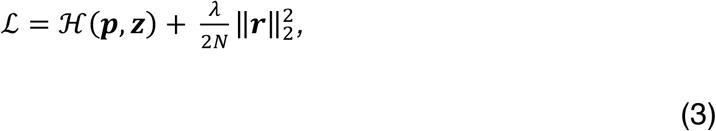

where the first part of eq. 3 denotes the cross-entropy loss *H*(**p**, ***z***) between the true probabilities of reward **p** (which were equal to 0 or 1, depending upon the stimuli for that trial) and the model’s readout probabilities ***z*** and the second term corresponds to a metabolic cost on all firing rates. Before training, the values of the **β**’s were drawn randomly from a Gaussian distribution with 0 mean and variance 0.05, and elements of **W**_out_ and ***b*** were set to 0. We trained two types of models: (1) with no regularisation (λ = 0; **Supp. Fig. 2a**) and (2) with high regularisation (λ = 20; **Supp. Fig. 2b**). In line with our predictions, selectivity to task variables in models trained with high regularisation converged on the minimal selectivity regime whereas models trained with no regularisation produced randomly mixed selectivity (**Supp. Fig. 2c,d**). This network formulation maps directly onto the selectivity space examined in experiment 2—stimulus set, stimulus set × context, and context—with context as the task-relevant variable as the XOR was not explicitly solved within the network but was provided as an input feature, differing from other inputs only in its relevance to the output.

### Analysis methods

#### Decoding

To test what information was represented by the observed neural population as a function of learning, we employed linear SVM decoding^7,14^. In contrast to the regression analysis, where the regression coefficients was estimated for every neuron separately, decoding analyses were run on pseudo-populations. This approach enhanced statistical power, reducing the likelihood of Type II errors, and mitigated the impact of session-specific sampling bias. Given the varying number of trials per session (both within and between animals), a sliding window method was employed to utilise all available data. Specifically, each session was divided into three windows, each matching the size of the session with the fewest trials (*n* = 801). Neural activities (*trials x neuron x time*) for the first 801 trials in each session during learning stage 1 were combined along the neuron axis to form a pseudo-population. This procedure was repeated for the middle and final 801 trials across each learning stage. This resulted in a matrix **X** = (X_*knt*_)_1≤*k*≤*K*,1≤*n*≤*N*,1≤*t*≤*T*_ for each of the time windows and each of the four learning stages, where the first dimension corresponds to K trials, the second dimension corresponds to N neurons (combined from two animals), and the last dimension corresponds to T time points. Then binary SVM classifiers were used to decode the task variables (colour, shape, width and XOR in experiment 1 and context, stimulus set, stimulus set x context (XOR 2), shape, width and XOR in experiment 2) at each time point for temporally resolved decoding (**Fig. 2e,f,g,i**; **Fig. 3e,f,g**; **Fig. 4a,e,i**; **Supp. Fig. 4i-l; Supp. Fig. 7a-c**). The statistical tests were run on firing rates average in broad time windows covering the entire feature presentation period (denoted by the pale orange shaded areas). An equivalent decoding procedure was used when analysing generative models (**Supp. Fig. 3e**). Decoding was performed in a cross-validated way where K trials were split randomly into set 1 and set 2, with each containing 50% of trials. The decoder was fitted using the set 1 and tested on set 2. The procedure was then repeated using set 2 as the training set and set 1 as the test set. Both decoding scores were then averaged. This procedure was repeated 10 times for different random splits of trials in sets 1 and 2, and these 10 resulting scores were then averaged. The decoding results were averaged over three time windows.

### Shattering dimensionality

To estimate shattering dimensionality^7,14^ in experiment 1(**Fig. 2j**), we used the same decoding approach as described above except that we averaged decoding scores over all 35 possible dichotomies that could be theoretically represented given the task structure (i.e., three linear variables form a cube in state space that can be dissected into two sets of 4 vertices in 35 possible ways; see **Supp. Fig. 10** for illustration). We estimated the decodability of each of these dichotomies and tracked their mean decoding accuracy as a function of learning. Exploring the theoretically maximal dimensionality of the colour-locked neural representations in experiment 2 using shattered dimensionality, where two linear variables (colour, stimulus set) were used, resulted in the identification of 3 (colour, stimulus set, colour x stimulus set (XOR2)) theoretically possible binary decoding problems. We tracked the mean of these dimensions over time (**Fig. 3h**).

### Cross-generalised decoding

To examine the neural geometry of the task variables we used cross-generalised SVM decoding^35^. In contrast to a typical cross-validation procedure, the testing happens not only on trials that were previously not seen but also on trials that correspond to different conditions. To achieve a high cross-generalisation score, it is therefore not sufficient to generalise across trial-wise noise but also to generalise across conditions^7^. Specifically, the labels of eight unique conditions (2 *colours x* 2 *shaps x* 2 *widhts* see **Supplementary Fig. 11a**) were split into two sets of four labels each, depending on the tested variable (e.g., colour 1 labels vs colour 2 labels; see **Supplementary Fig. 11b**). Each subset was further divided into a training set and a testing set (colour 1 vs colour 2 training set and colour 1 vs colour 2 testing set). A decoder was trained on the training set and then tested on the testing set, and vice versa; the two scores were then averaged. We identified 36 possible train-test splits (see **Supplementary Fig. 11c**), and the cross-generalised decoding score was obtained by averaging these scores. This method was used to determine whether the format of a task variable is abstract, meaning the variable is encoded in the same format as a function of the remaining task variables. Note that some of the train-test splits correspond to decoding a task variable as a function of a single other variable (e.g., decoding colour as a function of shape; see **Supplementary Fig. 11c**, shaded geometries), while others examine the decoding of the variable as a function of a mix of variables.

### Cross-stimulus set generalisation (decoding and selectivity analyses)

Cross-generalised decoding performed for experiment 2 data (both run on time-resolved and time-averaged firing rates; **Fig. 4a,e,i** and **Fig. 4b,f,j** respectively) differed in one aspect from the algorithm described in the *Cross-generalised decoding* section. As the aim of the analysis described here was to identify the neural format of the main task variables used across stimulus sets, only one splitting variable was used (i.e., stimulus set) to obtain cross-generalisation scores for the task-relevant variables (context, shape, and XOR). This reduced the possible cross-generalisation decoding axes to four possible binary decoding problems (e.g., when performing cross-generalised decoding for the colour variable we can: (1) train on differentiating colour 1 from colour 2 in stimulus set 1 and test on differentiating colour 3 from colour 4 in stimulus set 2, (2) train on differentiating colour 3 from colour 4 in stimulus set 2 and test on differentiating colour 1 from colour 2 in stimulus set 1, (3) train on differentiating colour 1 from colour 3 and test on differentiating colour 2 from colour 4, and (4) train on differentiating colour 2 from colour 4 and test on differentiating colour 1 from colour 3; these four decoding scores were then averaged). Using this procedure, we explored the cross-stimulus set generalisation potential of the colour, shape, width and XOR variables. Additionally, to test how selectivity of PFC cells changed as a function of learning in experiment 2 we employed a Pearson correlation metric. Specifically, we compared how similar the colour, shape and XOR coefficients in stimulus set 1 are to coefficients for the same variables in stimulus set 2 (**Fig. 4c, d, g, h, k, l**), which yielded three correlation scores for each of the main task variables (**Supp. Fig. 7d, e, f**). This was done for each of the four learning stages to explore whether selectivity for stimulus set 1 aligns with selectivity for stimulus set 2 as a function of learning, consistent with a shared abstract representation.

### Measuring similarity between selectivity distributions

To test the observed neural population for the presence of random mixed or minimal selectivity (**Fig. 1b,c**, and **Fig. 2k,l**), we firstly obtained regression coefficients for the three variables of interest (colour, shape and XOR; eq. 1) and constructed the selectivity space ***S***_*data*_. To assess the similarity of ***S***_*data*_ to ***S***_*minimal*_ and ***S***_*random*_, we computed the covariance matrix of ***S***_*data*_ (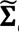_*data*_) as well as the covariance matrices of the expected random and minimal distributions given ***S***_*data*_ (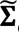_*minimal*_ and 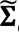_*random*_; see *Generative models)*. Finally, we calculated the normalised distance of the observed selectivity from model random selectivity

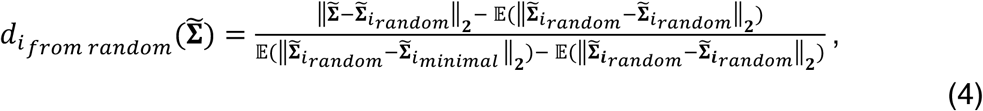

and the normalised distance of the observed selectivity to minimal selectivity

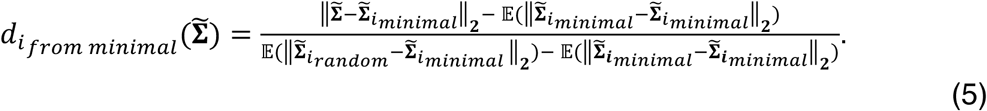

where the subscript *i* denotes a random draw and the expectations were computed over 1000 draws. From both the denominators and numerators, the distance within each of the models was subtracted to centre the measure around 0. More specifically, 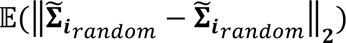 (the expected difference between two different randomly drawn selectivity distributions from the random model) was, for example, subtracted from the denominator and numerator of *d*_*from random*_ to account for within model distance. Additionally, both *d*_*from random*_ and *d*_*from minimal*_ were normalised by the distance between selectivities generated using both generative models 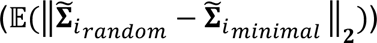 which resulted in the metrics being bounded between 0 and 1 (when 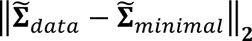 is equal or smaller than 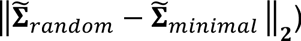. This was done to allow for a comparison of similarity estimates across learning stages. The Euclidean distance metric was chosen as the main analysis tool in this study based on simulations in which we generated different proportions of random and minimal selectivity across a single population (from 0% minimal and 100% of random to 100% minimal and 0% random) and compared the precision with which multiple metrics recovered the true proportions. We compared the Euclidean distance metric to the PAIRs metric, which has been used previously in the literature^12,28^, and to the symmetric Kullback–Leibler divergence estimate (KL divergence) which benefits from a strong theoretical basis and is assumption-agnostic. We found that, compared to the KL divergence and PAIRs metrics, the Euclidean distance measure yielded the highest precision of tracking learning-induced changes to neural selectivity (**Supp. Fig. 12a,b**). Specifically, our simulations showed that both the KL divergence and PAIRs can be used to precisely identify extreme selectivity regimes (either strong random selectivity or strong minimal selectivity) but fail at identifying intermediate selectivity regimes showing a strong bias towards random mixed selectivity (**Supp. Fig. 12a,b**). As the focus of this study was to track learning dynamics, a metric that allows to identify a broad range of selectivity regimes was chosen for the final analysis. Nonetheless, the results from experiment 1 (**Fig. 2m,n**) were broadly replicated using the symmetric KL divergence estimate (**Supp. Fig. 12c,d**) and PAIRs (**Supp. Fig. 12e,f**).

### Principal Component Analysis

PCA was used as a measure of neural dimensionality in experiment 1(**Supp. Fig. 4h**). Firstly, pseudo populations were constructed for each learning stage using the same procedure as described in the *Decoding* section. Then, firing rates were averaged in the time window preceding the outcome presentation ([t_400*ms*_, t_500*ms*_], shape-locked;). Next, principal components were run on condition averages. This was done separately for each learning stage. To compute how the variance explained (ratio) by the first PC changed as a function of learning, trials were randomly split into test and train 10 times; PCA was fitted then on train trials and the test trial firing rates were projected onto them to compute variance explained. The results from 10 random splits were then averaged. Note that width 1 and width 2 trials were pooled together. The null distribution for the permutation test was computed by randomly shuffling neurons between stage 1 and stage 4, and repeating the described PCA procedure (*n* = 500).

### Statistical testing

#### Decoding and cross. gen. decoding

Throughout the study, we employed non-parametric permutation tests to test statistical significance within each learning stage and between learning stages (learning-induced effects). Two types of null distributions were thus constructed: (1) for statistical testing of above chance-level decoding analyses the labels describing the trial dimension (*k*) of the pseudo-population matrix **X** = (X_*knt*_)_1≤*k*≤*K*,1≤*n*≤*N*,1≤*t*≤*T*_ were randomly permuted 1000 times; (2) to test for learning-induced effects on decoding scores, the matrices **X**_*stage* 1_ and **X**_*stage* 4_ were concatenated along the neuron dimension (*n*) and then 1000 new **X**′_*stage* 1_ and **X**′_*stage* 4_ matrices were generated by randomly assigning neurons to either **X**′_*stage* 1_ or **X**′_*stage* 4_. One-sided tests were used when testing the predictions of the minimal model and two-sided tests were used when no differences were expected.

#### Selectivity measures

To test whether observed selectivity was dissimilar to the random selectivity regime and similar to minimal selectivity regime 1000 random and minimal models were generated using data-derived parameters for each learning stage. Next, the *d*_*rand.from rand*_. and *d*_*rand.from min*_. distances were computed for 1000 randomly generated models according to eq. 4 and eq. 5 (with 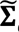_*random*_ as input) to serve as null distributions for both comparisons. Note that the observed selectivity was compared to random model selectivity when analysing the data’s similarity to random (**Fig. 2m**; **Fig. 3m**) as well as minimal selectivity (**Fig. 2n**, **Fig. 3n**). Furthermore, as in experiment 2 we tested whether selectivity for task variables was similar in stimulus set 1 to variables in stimulus set 2 and whether this selectivity alignment changed over learning, two null distributions were thus constructed: (1) statistically significant selectivity alignment was assessed by comparing the observed correlation to a distribution (*n* = 1000) of correlations obtained after shuffling one of the selectivity vectors; (2) learning-induced effects in selectivity alignment were assessed by comparing the observed difference in alignment between stage 1 and stage 4 to a distribution of differences computed after randomly shuffling neurons between stage 1 and 4.

## Supplementary materials

### Section 1.1: How to set selectivity parameters for optimal XOR decoding

We assume that neural activities ***x*** of ***N*** neurons are given by the following regression model

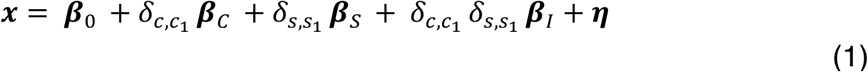

Where **β**_*C*_, **β**_*S*_, **β**_1_ are the regression ‘coefficients’ for colour, shape, and the interaction term, respectively, δ_*c*,*c*2_ = 1 if the colour *c* is colour 1 and -1 otherwise (same for δ_*s*,*s*2_ for shapes *s*), and ***η***∼ *N*(**2**, Σ_1_). Therefore,

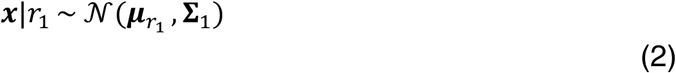

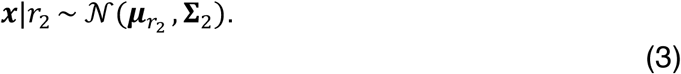

Where *r*_1_ is one XOR condition (i.e., *c* = 1, *s* = 1 or *c* = 2, *s* = 2) and *r*_2_ is the other XOR condition (i.e., *c* = 1, *s* = 2 or *c* = 2, *s* = 1) and Σ_2_ is some noise covariance matrix. Note that with the inclusion of the interaction term, it is sufficient to separate the two XOR conditions. We now calculate μ_*r*2_ and μ_*r*3_ :

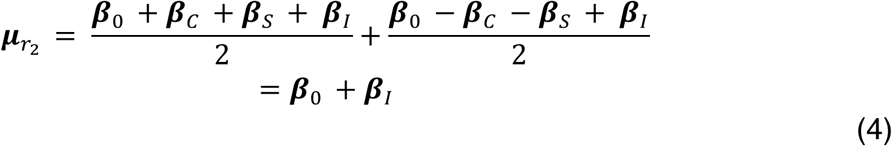

and,

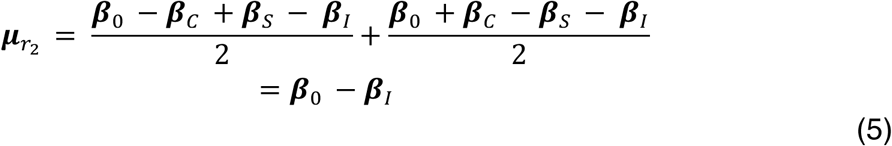

Therefore,

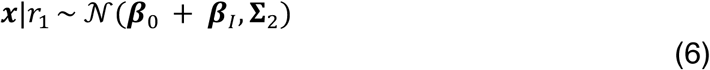

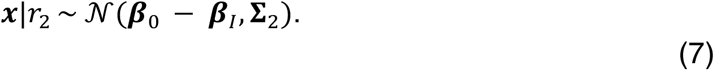

Therefore, we need **β**_1_ > 0 to be able to separate the two XOR conditions.

### Section 1.2: An energy cost on unnecessary neural activity

If we also consider minimising the squared norm of neural activity for each condition (i.e., an energy cost), we have

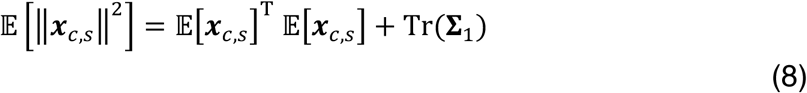

where the subscripts *c* and *s* correspond to colour and shape indices, respectively. Therefore,

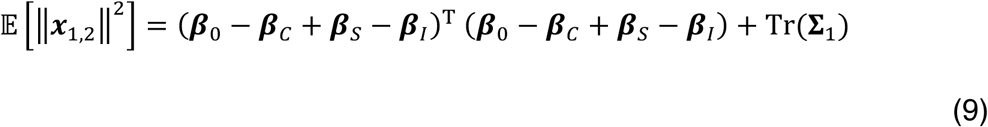

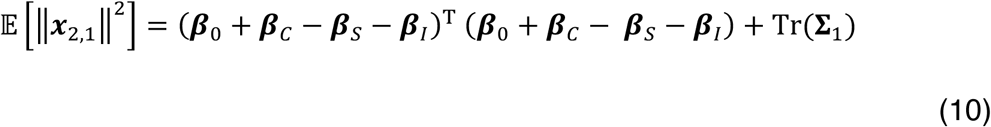

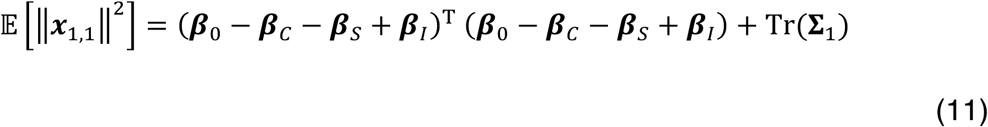

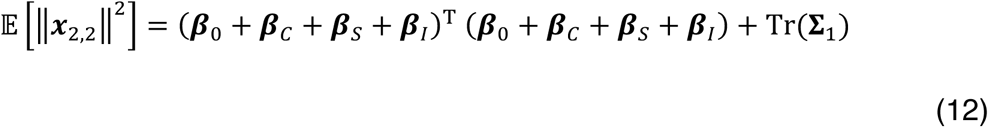

Therefore, the total mean energy cost *m* is given by

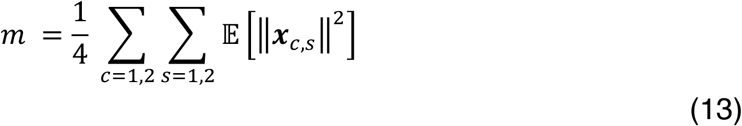

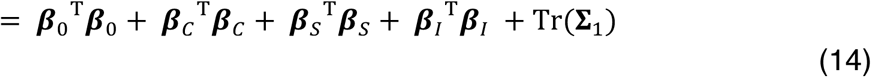

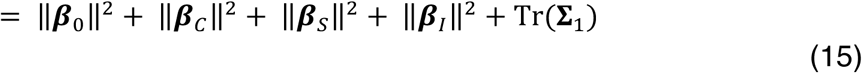

To minimise *m* while keeping **β**_1_ > 0, which we need for performance, we can set **β**_0_ = **β**_*C*_ = **β**_*S*_ = **2** which gives

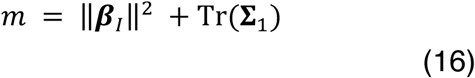

## Section 2: supplementary figures

**Supplementary figure 1.**
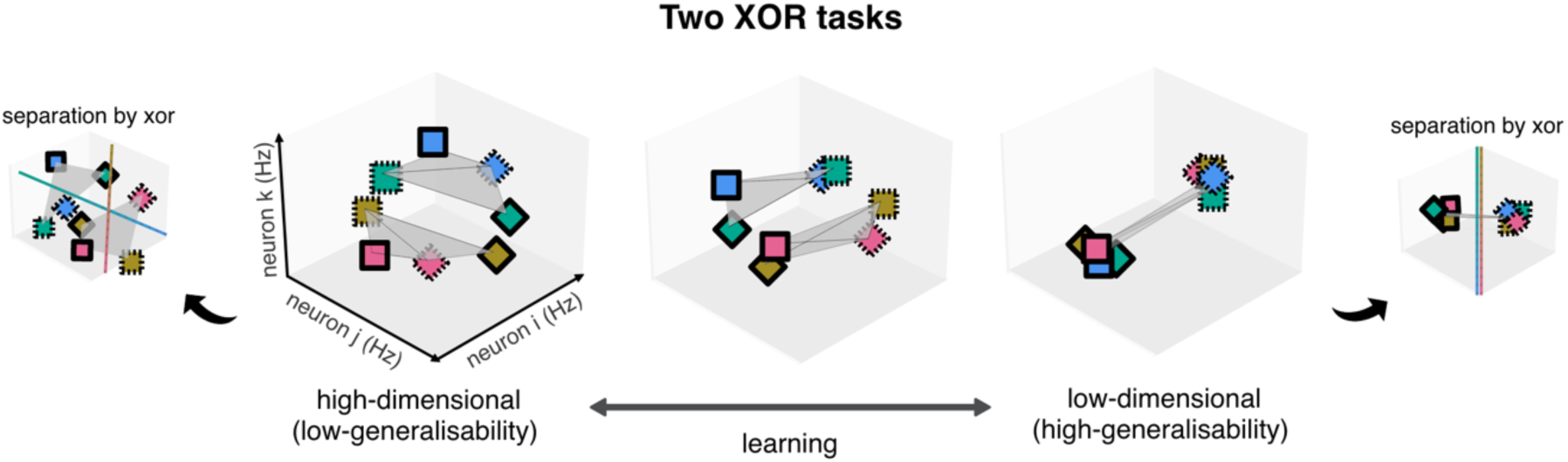
Different solutions to the discriminability-generalisability trade-off when a new stimulus set is being learnt. Low-dimensional representations are more likely to support high generalisability. When the task representation is high-dimensional (left), aligning new with old stimuli becomes a harder problem as all task variables must be jointly aligned; in this case, it is not trivial that the XOR discriminant from one task would correctly differentiate the XOR feature in the second task, particularly if high discriminability of the remaining variables is to be maintained. By contrast, when the neural code is low-dimensional (right), both tasks can more readily be aligned to a common axis, enabling a shared XOR discriminant that generalises across tasks.

**Supplementary figure 2.**
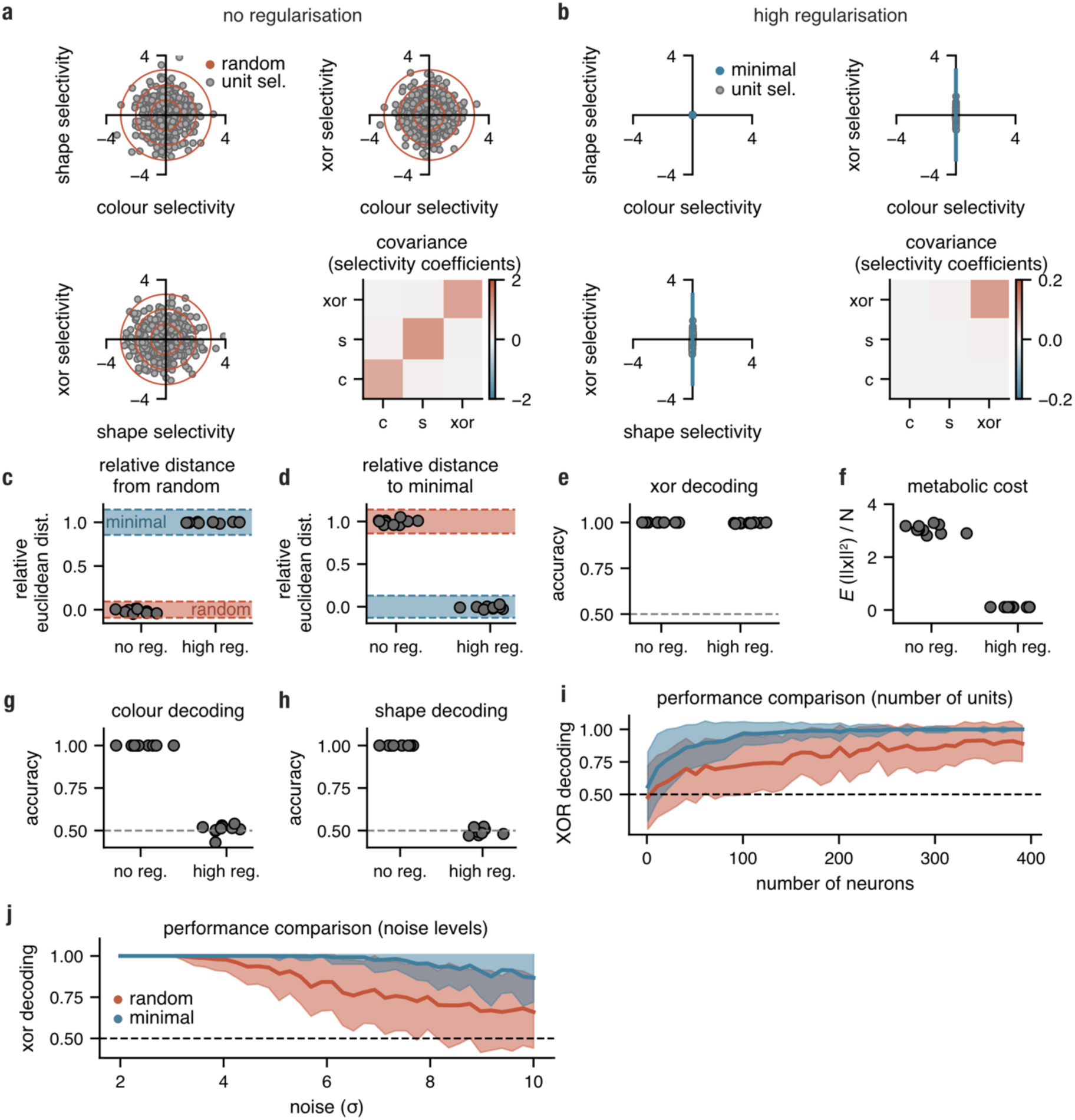
Optimised feedforward networks converge to the minimal XOR selectivity model. Twenty feedforward networks were trained (10 with high levels of regularisation and 10 with no regularisation) to perform the XOR task. **a, b,** Selectivity observed in no and high regularisation networks, respectively; models of random (red ellipses) and minimal selectivity (blue ellipses) well approximated the observed selectivity. **c, d**, After training, low regularisation networks converged on a random mixed selectivity regime and high regularisation networks on a minimal XOR regime. **e**, post-training XOR decoding (linear SVM) for both no and high regularisation models. **f**, No regularisation models exhibited substantially lower metabolic cost (cf. **Supp. materials eq. 8**). **g**, **h**, Colour and shape decoding (linear SVM) for no and high regularisation models, respectively. **i**, **j**, Comparison of XOR decoding obtained from minimal and random generative models as a function of population size (i) and noise (σ) levels (j). Dashed grey line shows chance-level decoding.

**Supplementary figure 3.**
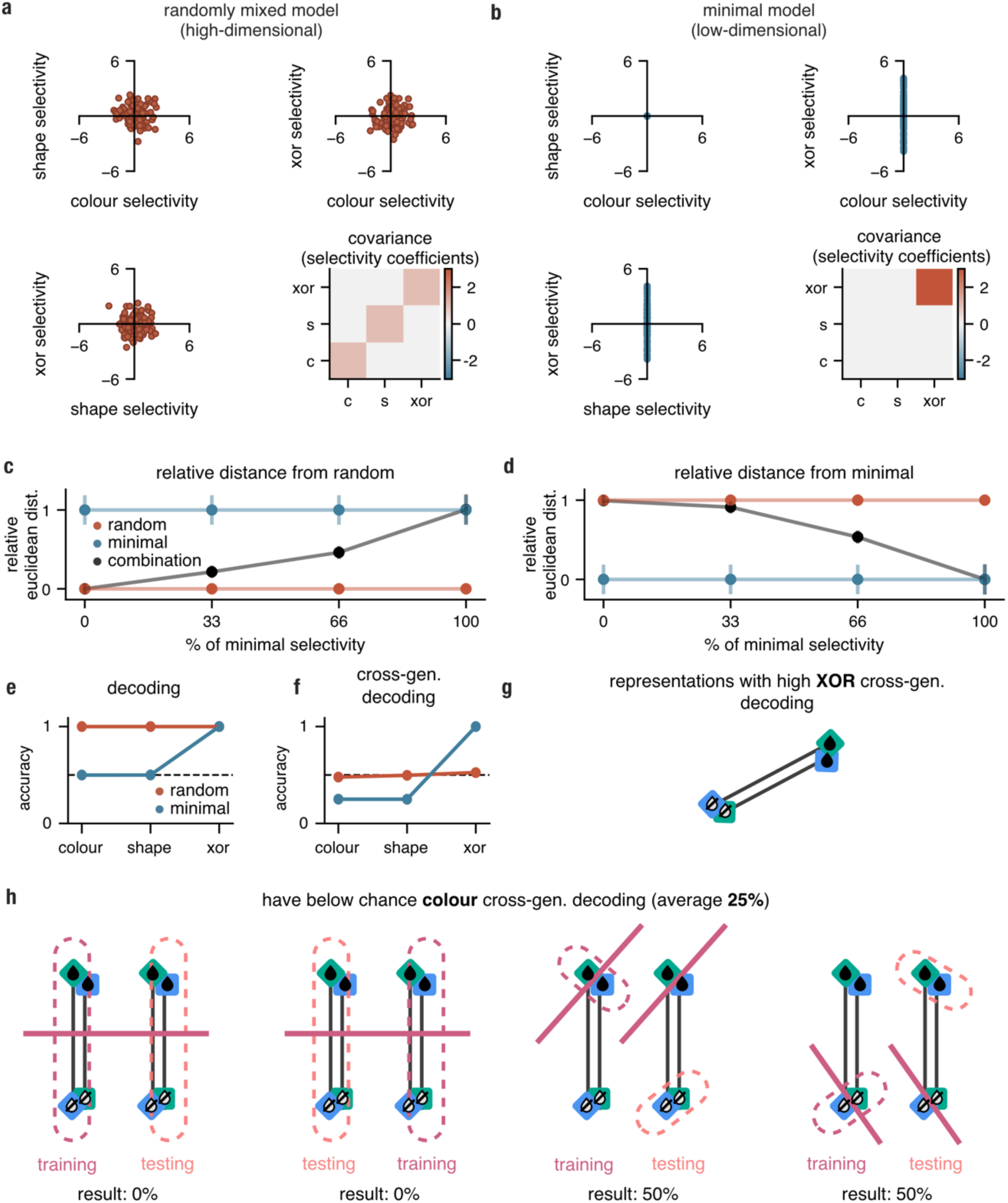
Predictions from random mixed and minimal selectivity models. **a**, Each neuron can be represented as a point in the 3-dimensional selectivity space spanned by colour, shape, and XOR (their interaction). In the random mixed model, selectivity is distributed according to a spherical Gaussian distribution in this space (Methods, generative models); the covariance matrix is computed between the selectivity coefficients. **b**, Analogous to a but for the minimal model; neurons are selective only for the XOR (interaction between colour and shape), as this is the only feature that is necessary to solve the task. **c**, Relative distance between the covariance matrices of either the random (red), minimal (blue) or the model with varying proportions of minimal selectivity (black) to the random selectivity model. **d**, Same as panel c but the distance is calculated relative to the minimal selectivity model. Red and blue error bars show standard deviations (±1 *s. d*. over 1000 randomly drawn models; see *Methods*, *measuring similarity between selectivity distributions*) of the relative Euclidean distance between the covariance matrix of the random and the minimal model (with matched total variance to the data; standard deviation of random-to-minimal distance is too small to be visible; red) and relative Euclidean distance between the covariance matrix expected from two randomly drawn minimal models (blue); black error bars show the standard deviation of the relative distance between the surrogate covariance (with varying proportions of minimal selectivity) and random covariance (±1 *s. d*. over 1000 random models). **e**, Mean (over 100 models) decoding of task variables for the random (red) and minimal (blue) models. Dashed grey line shows chance-level decoding. **f**, Mean (over 100 models) cross-generalised decoding of task variables for the random (red) and minimal (blue) models. Dashed grey line shows chance-level decoding. **g**, **h**, Relationship between high XOR cross-gen. decoding and below chance colour cross-gen. decoding. The clustering of all rewarded trials (XOR == True) and non-rewarded trials (XOR == False) on opposite sides of an axis (representing the abstract XOR) results in high cross-gen. decoding for the XOR (g). Consequently, colour exhibits below-chance cross-gen. decoding. This occurs because some colour decoding axes are inverted between training and testing splits, leading to 0% accuracy scores (h; left), while others assign a single class label to all examples when testing, resulting in a 50% score (h; right). The same argument holds for shape.

**Supplementary figure 4.**
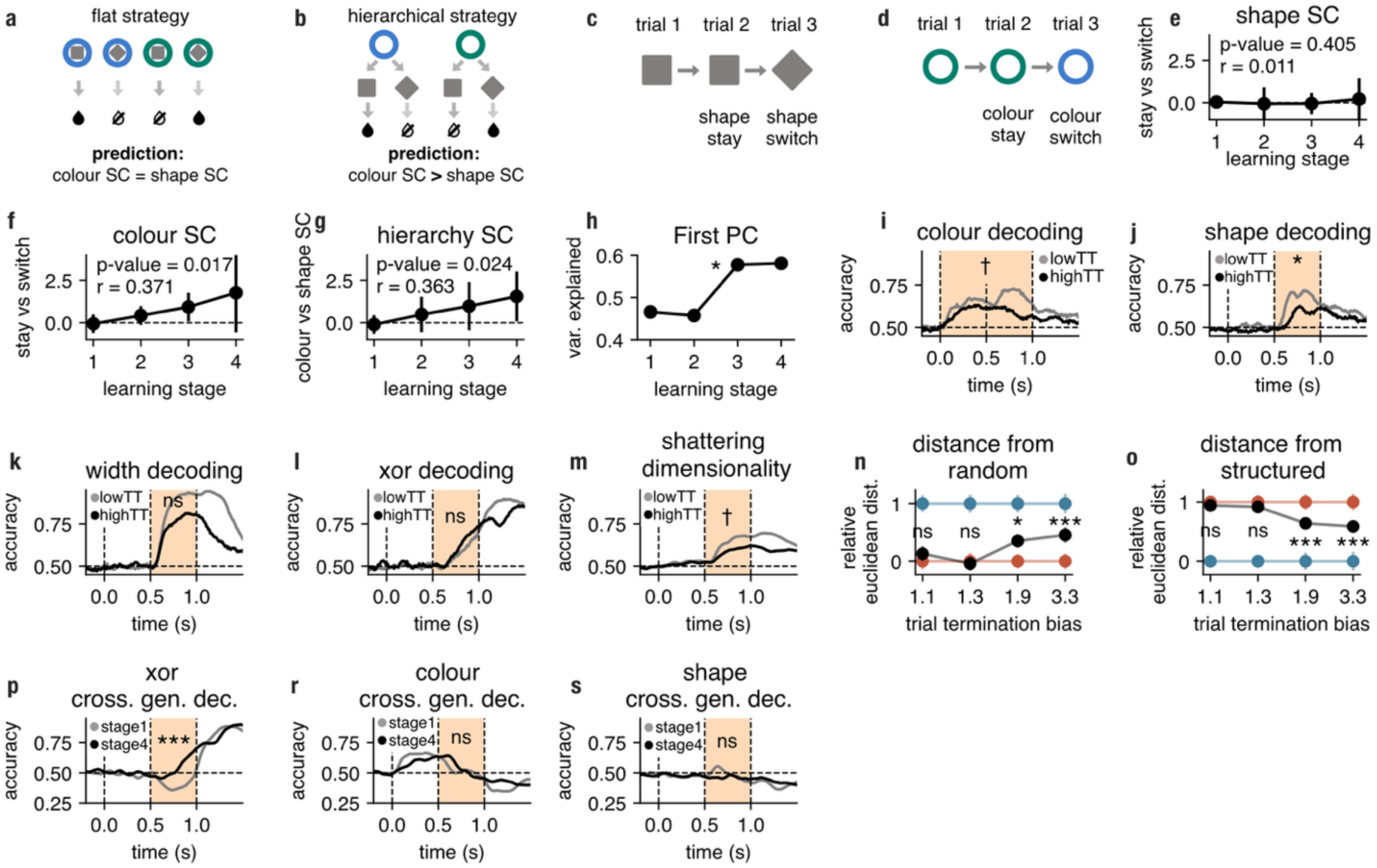
Decoding of task variables as a function of learning and behaviour in experiment 1. **a, b**, The animals could adopt two possible strategies to learn the task: (a) memorising every stimulus–outcome pairing, or (b) using a hierarchical strategy in which colour served as a first-order policy cue, guiding subsequent context-dependent processing of shape. To differentiate between these strategies, colour and shape switch costs were computed. **c**, Shape switch costs: trial termination rates were compared between trials in which the shape changed vs. remained the same on consecutive trials. **d**, Analogous illustration but for colour switch costs. **e, f, g**, Trial termination switch costs for colour, shape, and the difference between colour switch costs and shape switch costs, plotted as a function of learning stage. **h**, variance explained (ratio) by the first principal component plotted as a function of learning (see *Methods, principal component analysis* for details). **i-o,** Analogous to Figure 2 but run on sessions sorted by proportion of adaptive trial termination (TT); statistical tests were conducted on firing rates averaged over the time window indicated in pale orange. **p, r, s**, Temporally resolved cross-generalised decoding of XOR(p), colour(r), and shape(s) in stage 1 and stage 4; the pale orange shaded areas indicate the time window in which statistical tests were run. All p-values were calculated from permutation tests (***, *p* < 0.01; **, *p* < 0.01; *, *p* < 0.05; †, *p* < 0.01; *n. s*., not significant).

**Supplementary figure 5.**
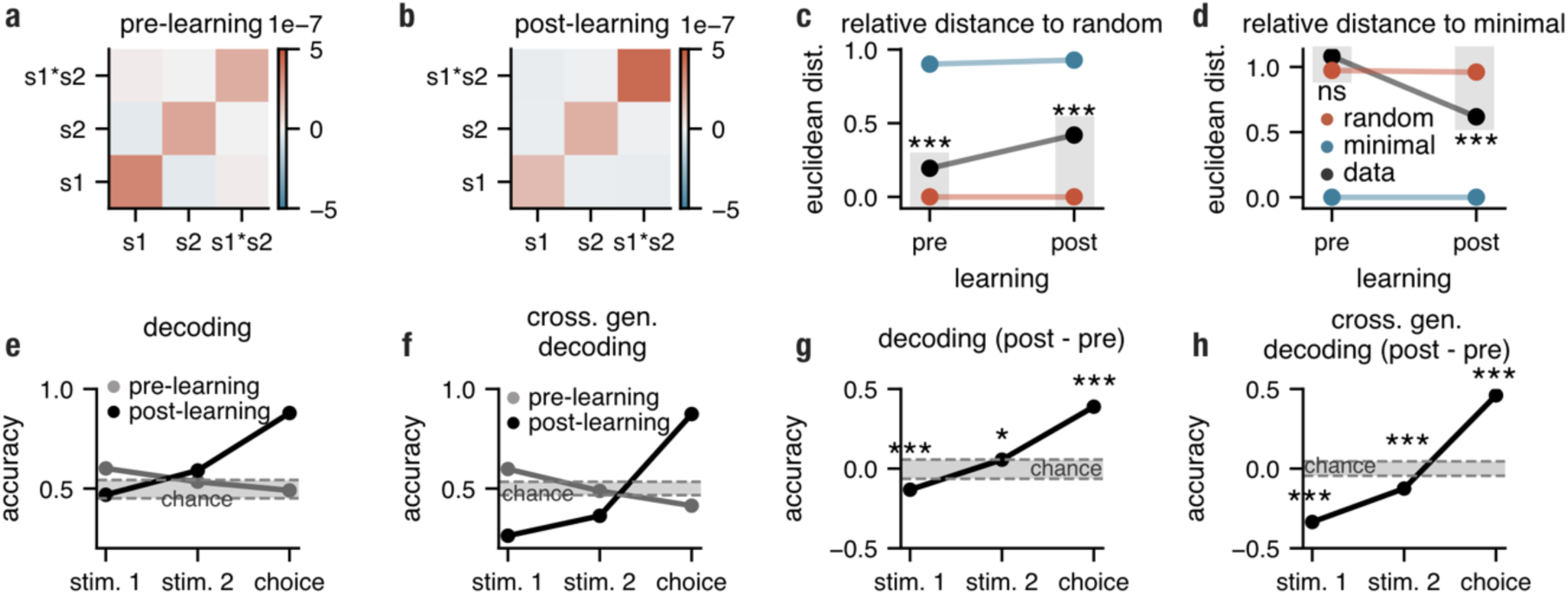
The re-analysis of Constantinidis et al.^25–27^ dataset. We employed the same analysis methods as in Fig. 2 to test whether the PFC activity reported in Constantinidis et al. converged on a minimal XOR model. **a**, the covariance matrix describing relations between the selectivity for stimulus 1, stimulus 2 and their interaction (XOR) in the pre-learning phase of the experiment. **b**, Same as panel a but post learning. **c**, Relative Euclidean distance between the covariance matrix of observed selectivity coefficients over learning and the covariance matrix expected from random selectivity (with matched total variance) (*Methods*, *measuring similarity between selectivity distributions*). **d**, Same as panel **c** but we show the relative distance from the covariance matrix expected from minimal selectivity (with matched total variance). **e**, Decoding of task variables for pre- and post-learning stages. **f**, Cross-generalised decoding of task variables plotted as a function of learning. **g, h**, Learning-induced accuracy differences in decoding and cross-generalised decoding, respectively. Shaded areas in **e-h** illustrate chance-level decoding obtained by shuffling trial labels (for details see *Methods, statistical testing*). All p-values were calculated from permutation tests (***, p < 0.01; **, p <0.01; *, p < 0.05; n.s., not significant).

**Supplementary figure 6.**
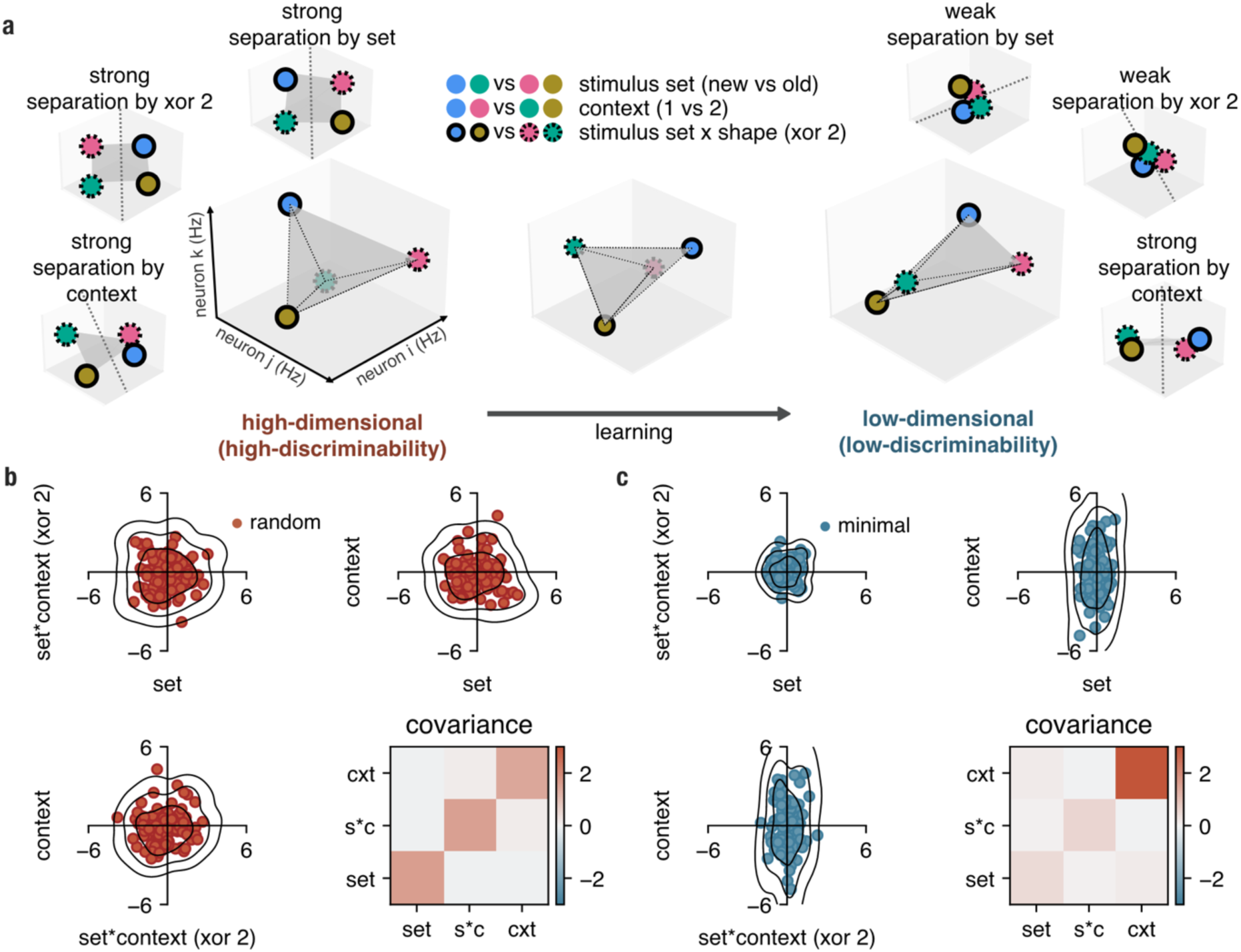
Hypothesised effects of learning on neural geometry and selectivity in the prefrontal cortex in experiment 2. Like in Figure 1, learning is depicted as reducing neural dimensionality, changing how many strong linear decoding axes can be implemented on neural firing rates (discriminability). **a**, A high-dimensional regime allows the strong separation of all task features using three possible readout axes (left), whereas a low-dimensional representation only allows task-relevant features to be strongly separated (right). **b**, Each neuron can be represented as a point in the 3-dimensional selectivity space spanned by stimulus set, stimulus set x context (XOR 2), and context (Fig. 3b). In the random model, selectivity is distributed according to a spherical Gaussian distribution in this space (Methods, generative models); the covariance matrix is computed between the selectivity coefficients; zero-mean Gaussian noise (σ = 0.7) was added to each selectivity coefficient to illustrate measurement bias under finite sampling. **c**, Analogous to b but for the minimal model; neurons are strongly selective only for the context, as this is the only feature that is necessary to solve the task.

**Supplementary figure 7.**
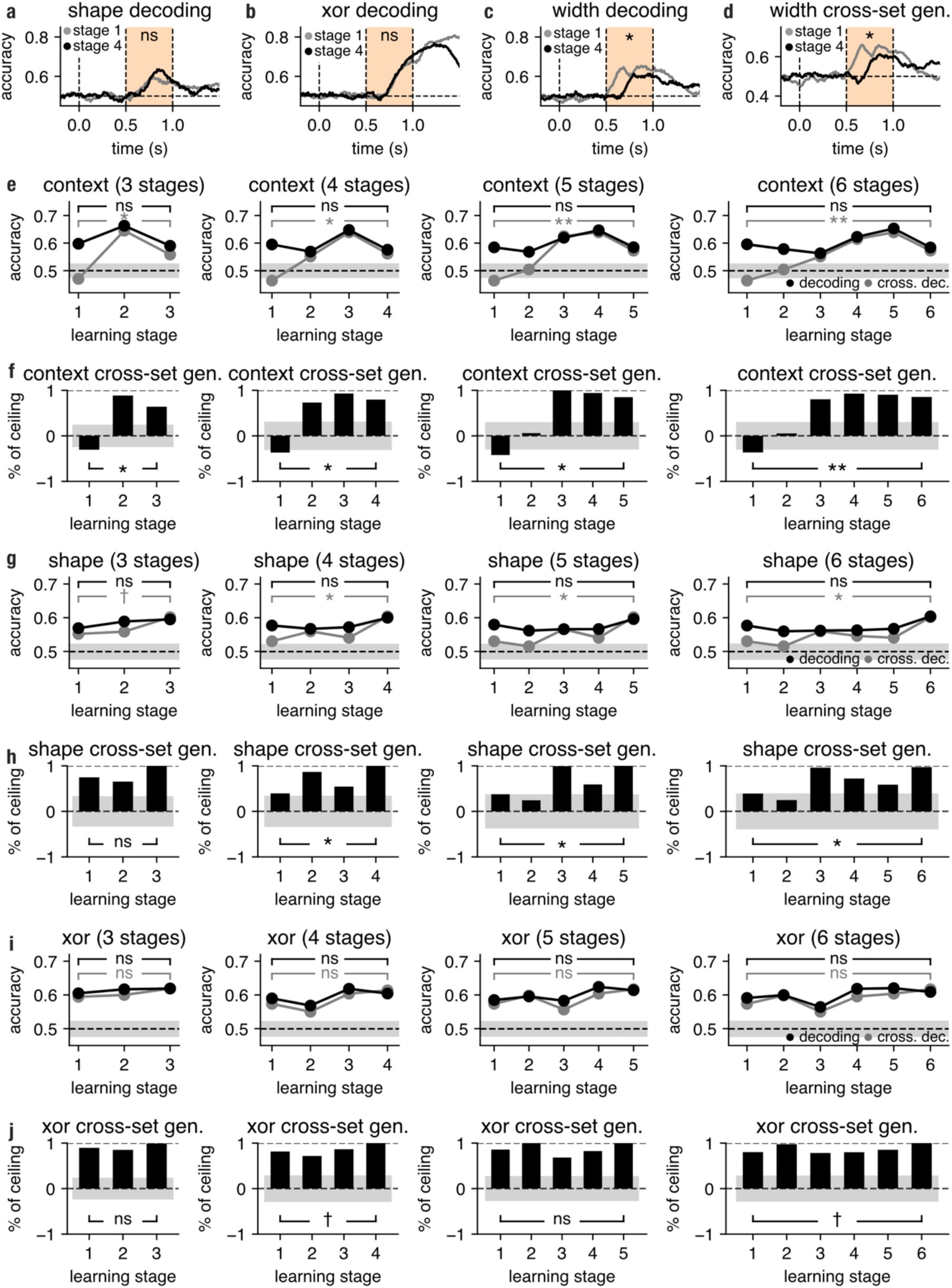
Decoding and selectivity results in experiment 2. **a-c**, Temporally resolved linear SVM decoding for shape, XOR and width; the pale orange shaded areas indicate the time window in which statistical tests were run. Horizontal dotted lines represent chance-level decoding whereas vertical dotted lines indicate the onset of the colour, shape and the trial outcome. **d**, Temporally resolved cross-generalised decoding of width (trained on set 1 and tested on set 2 (and vice-versa)). **e**, Context cross-set generalisation results remain consistent across different learning discretisations; analogues to Figure 4d computed for 3, 4 (original), 5 and 6 learning stages (see *Methods*, *data acquisition and pre-processing* for details). **f**, Cross-set context generalisation normalised by linear context decoding (cf. panel e, black vs grey lines). Cross-set generalisation performance is expressed as a proportion of linear context decoding (ceiling) across learning stages. Positive values indicate geometries in which colour 1 vs colour 2 generalises to colour 3 vs colour 4, whereas negative values indicate the flipped geometry (colour 1 vs colour 2 generalises to colour 4 vs colour 3). Grey shaded areas indicate the null distribution obtained by shuffling trial labels. **g**,**h**, Analogous to e-f but computed for shape cross-set generalisation. **i**,**j**, Analogous to e-f but computed for XOR cross-set generalisation. All p-values were calculated from permutation tests (***, *p* < 0.01; **, *p* < 0.01; *, *p* < 0.05; †, *p* < 0.01; *n. s*., not significant).

**Supplementary figure 8.**
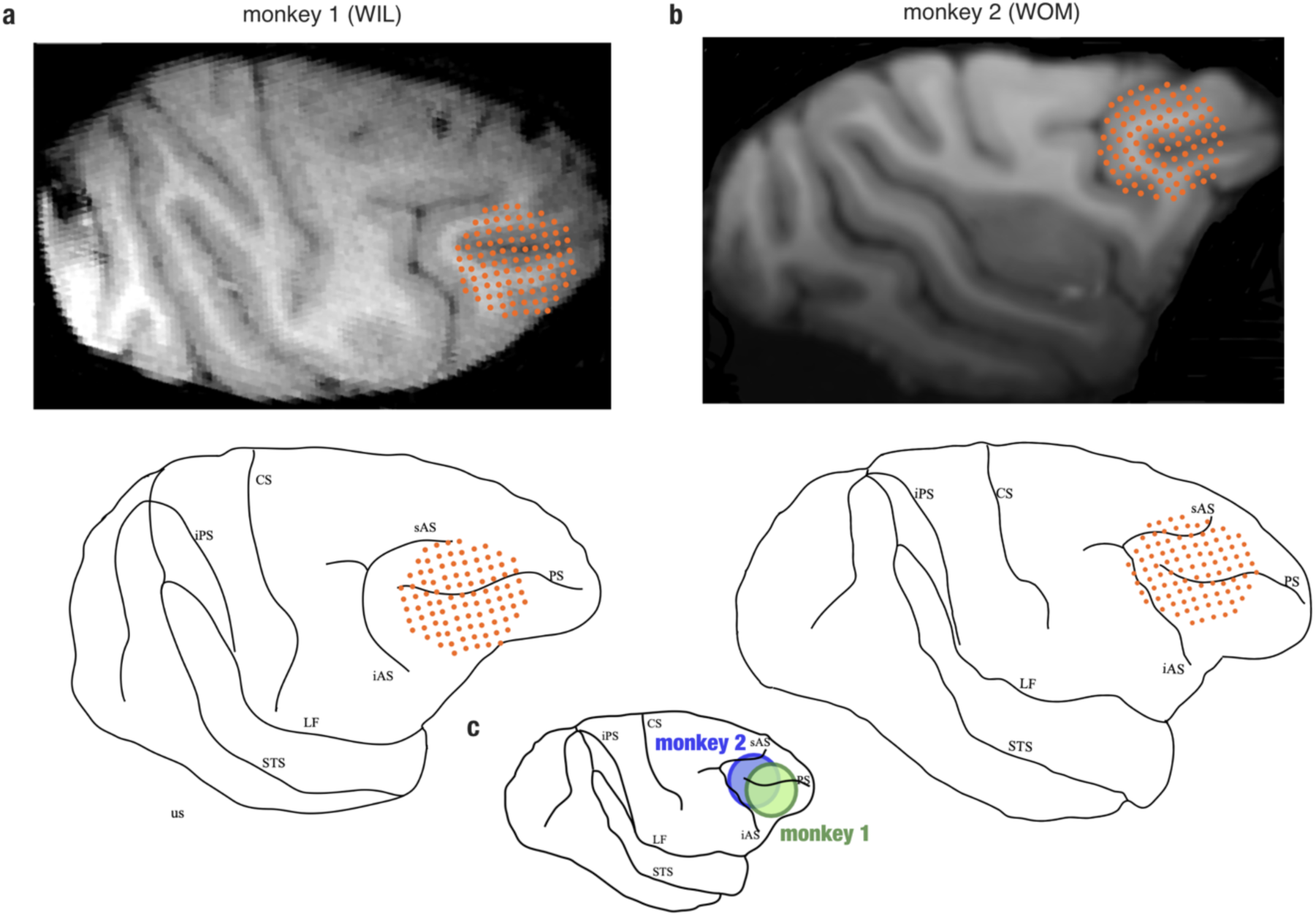
Electrode locations in monkey 1 (a) and monkey 2 (b) and their comparison (c).

**Supplementary figure 9.**
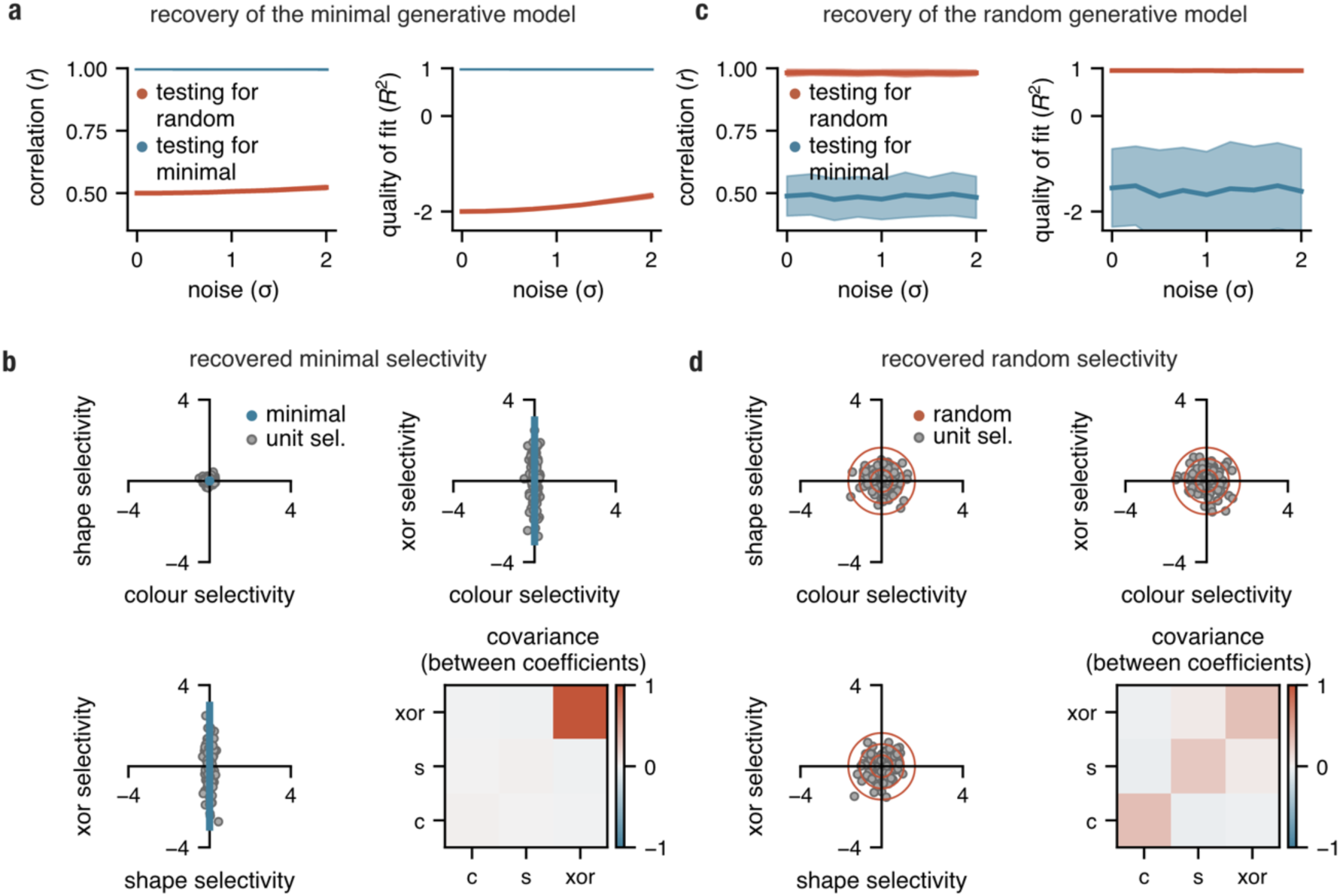
Linear regression recovers underlying generative models. **a**, Pearson correlation and R^2^ computed between the covariance matrix of recovered selectivity and true underlying selectivity (when minimal generative model was used) for different levels of noise. **b**, Selectivity coefficients obtained after running a linear regression plotted for each unit in selectivity space (for σ = 2); minimal model overlaid in blue; covariance matrix computed between the recovered selectivity coefficients. **c**,**d**, Analogous to a,b but when the random model was used to generate data; random model overlaid in red. Shaded areas in a and c indicate the mean ±1 *s. d*. computed over 100 different initialisations.

**Supplementary figure 10.**
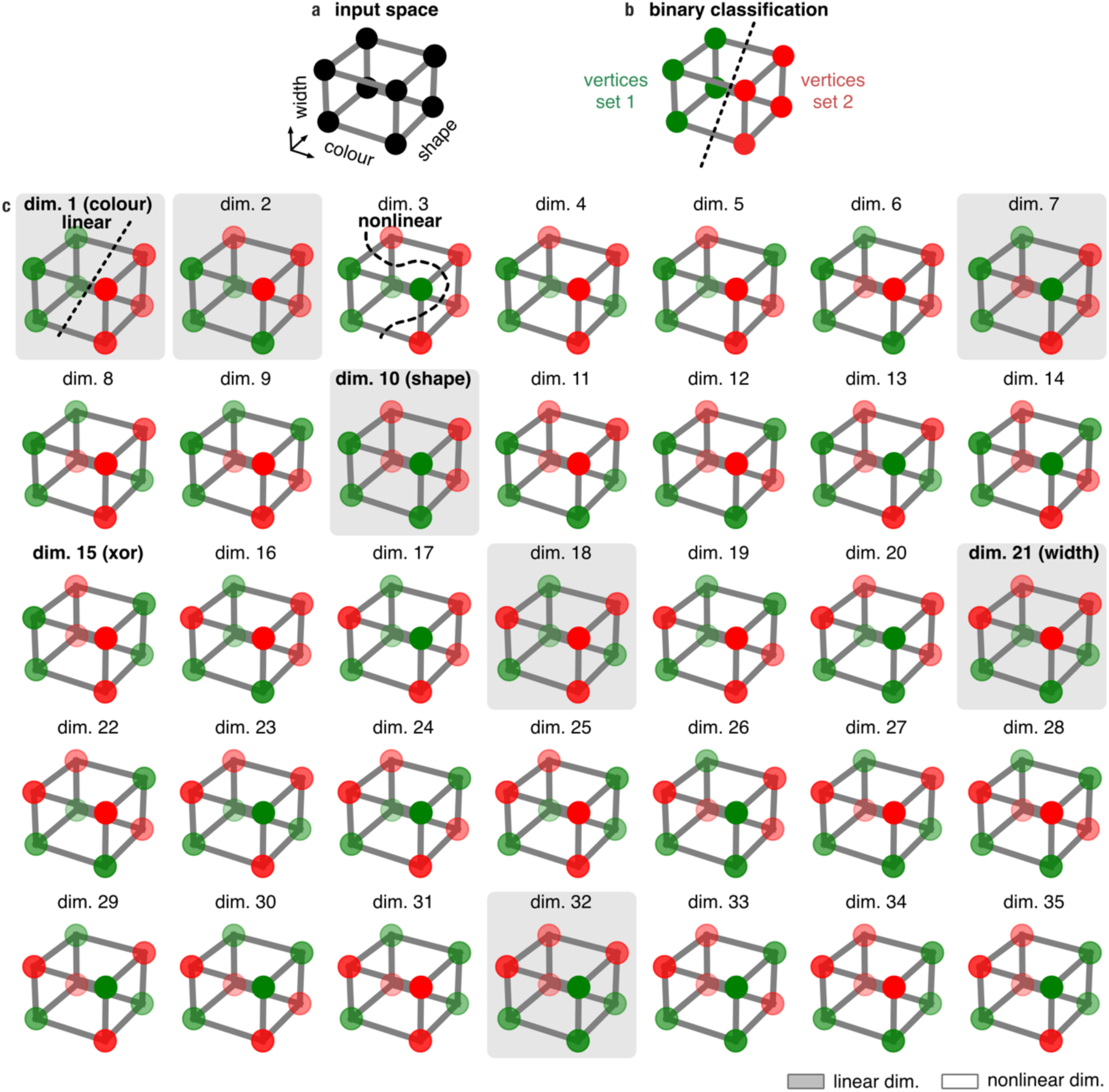
Schematic depiction of shattering dimensionality in the XOR task in experiment 1. **a**. Colour, shape, and width form a cube in the input space, with each of the 8 vertices representing a unique combination of these input variables. **b**. The vertices can be divided into two equally sized groups of four vertices each, representing a binary classification problem. **c**. All 35 theoretically possible binary problems (red vs. green vertices) are obtained by randomly splitting the vertices into two equally sized groups. Dimensions 1, 10, 15, and 21 correspond to colour, shape, XOR, and width, respectively. The grey and white backgrounds indicate whether a dimension is linear (can be split by a plane, e.g. dimension 1) or nonlinear (cannot be split by a plane, e.g. dimension 3), respectively. Colour intensity corresponds to spatial depth, with solid colours signifying vertices closer to the viewer and pale colours signifying those farther away.

**Supplementary figure 11.**
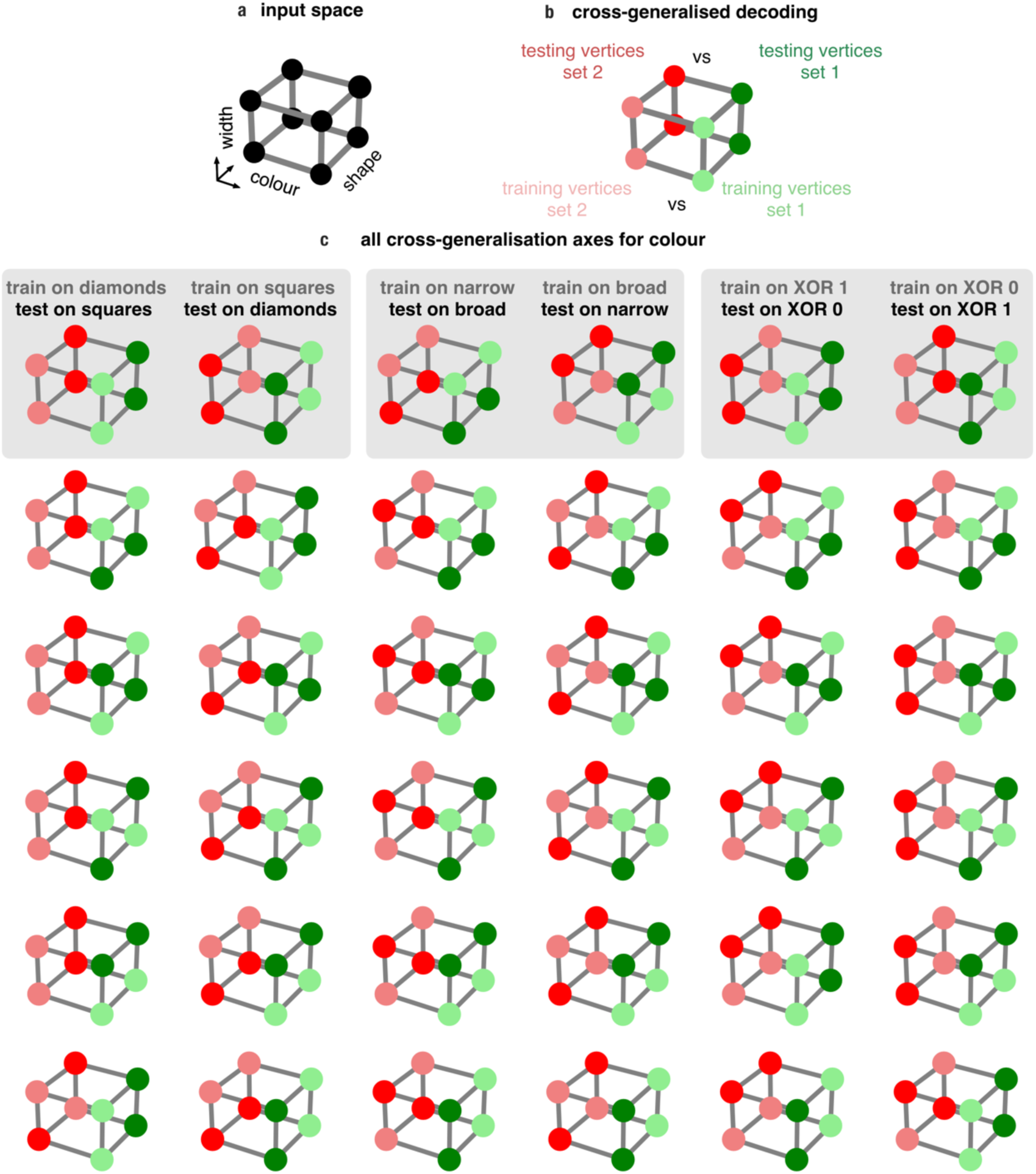
Schematic depiction of cross-generalised decoding of the variables. **a**. Colour (blue vs green), shape (diamond vs square), and width (narrow vs broad) form a cube in the input space, with each of the 8 vertices representing a unique combination of these input variables. **b**, To test whether a variable is encoded in an abstract format relative to a single variable, two binary linear classification problem were defined. Firstly, a classifier was trained on differentiating that variable (e.g. colour) only on a subset of trials (e.g., diamond shape trials). Next, this classifier was tested on the remaining subset of trials (e.g., square shape trials). For example, a high score obtained from such a decoding procedure indicates that the same representation of colour was used as a function of different levels of shape, a hallmark of abstract coding. **c**, To test whether a variable (e.g., colour) has an abstract format relative to all remaining task variables, this procedure needs to be repeated for all possible train and test splits (when colour 1 is always on the left and colour 2 is always on the right side of the cube). These 36 scores are then averaged to obtain cross-generalised decoding of colour. Some of the test-train splits correspond to task variables like shape, width and XOR (grey background) while others represent a mixture of input variables.

**Supplementary figure 12.**
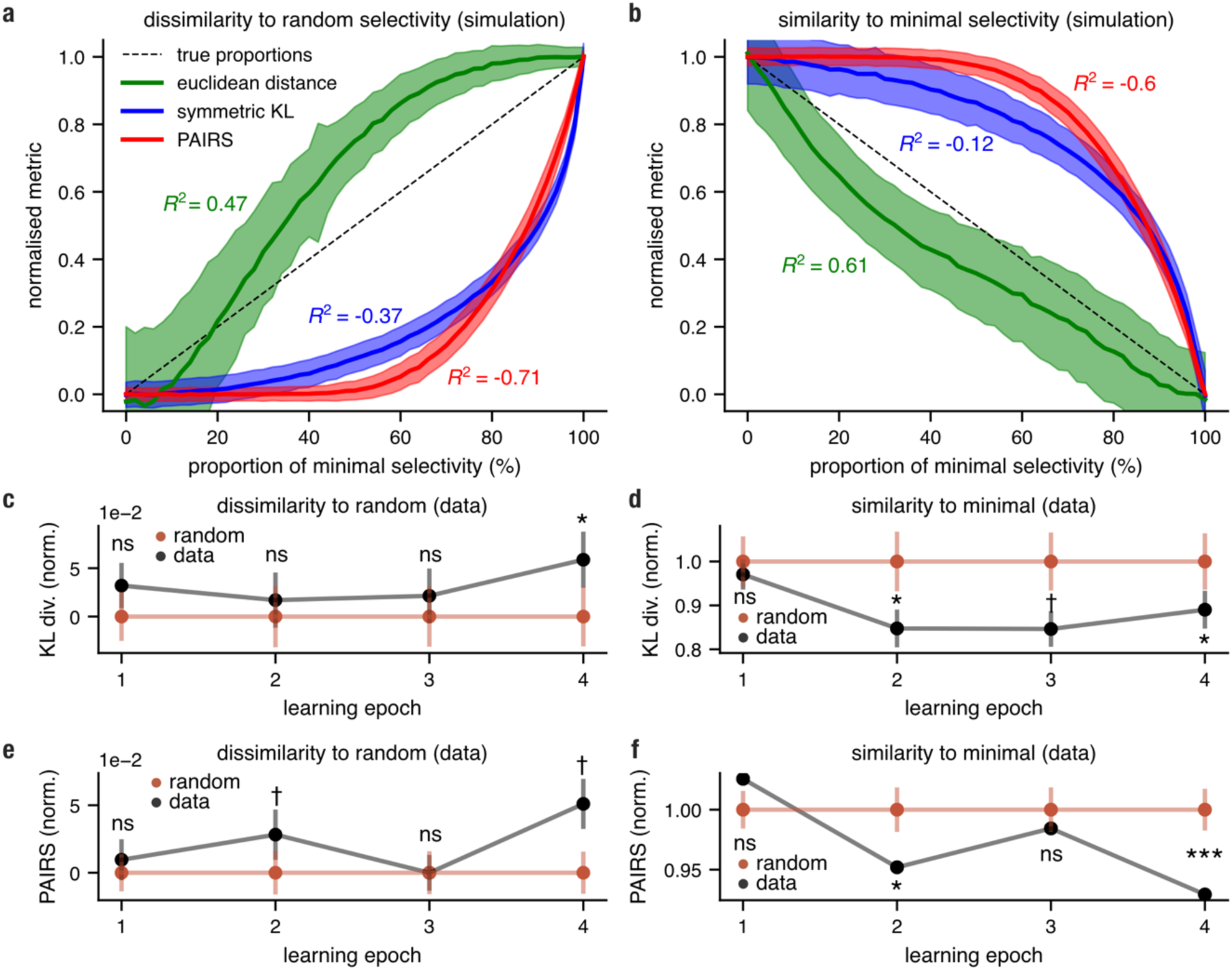
The precision of Euclidean distance, symmetric KL estimate and PAIRS metrics in tracking learning-induced changes to neural selectivity. **a**, Relative Euclidean distance between the covariance matrix of selectivity coefficients obtained from a simulated mixed population (random-minimal) and the covariance matrix expected from pure random selectivity plotted as a function of minimal selectivity proportions (0-100%; for details see *methods*, *measuring similarity between selectivity distributions*). Coloured annotations indicate mean R^2^ values computed between the true proportions (dotted lines) and estimated proportions (coloured bold lines). **b**, Same as panel **a** but we show the relative distance from the covariance matrix expected from minimal selectivity; shaded areas illustrate mean ±1 *s. d*. for each of the metrics computed from 1000 randomly drawn selectivity models. **c, d** Selectivity results from experiment 1(**Fig. 2m,n**) computed using symmetric KL divergence estimate. **e, f** Selectivity results from experiment 1(**Fig. 2m,n**) computed using the PAIRS metric.

## Notes

https://datadryad.org/stash/dataset/doi:10.5061/dryad.c2fqz61kb

### Competing Interest Statement

The authors have declared no competing interest.

### Summary of Updates

Figure 4 and Supp. Figure 7 were updated to show that the cross-set generalisation effect is robust to analysis window and learning discretisation decisions.

